# Synchronous multi-segmental activity between metachronal waves controls locomotion speed in *Drosophila* larvae

**DOI:** 10.1101/2022.09.08.507222

**Authors:** Yingtao Liu, Eri Hasegawa, Akinao Nose, Maarten F. Zwart, Hiroshi Kohsaka

**Affiliations:** Department of Physics, Graduate School of Science, the University of Tokyo, 7-3-1 Hongo, Bunkyo-ku, 133-0033 Tokyo, Japan; Department of Complexity Science and Engineering, Graduate School of Frontier Science, the University of Tokyo, 5-1-5 Kashiwanoha, Kashiwa, 277-8561 Chiba, Japan; School of Psychology and Neuroscience, Centre of Biophotonics, University of St Andrews, St Andrews KY16 9JP, UK; Graduate School of Informatics and Engineering, the University of Electro-Communications, 1-5-1, Chofugaoka, Chofu-shi, 182-8585 Tokyo, Japan; Frontier Science and Social Co-creation Initiative, Kanazawa University, Kakuma, Kanazawa, 920-1192, Ishikawa, Japan

## Abstract

The ability to adjust the speed of locomotion is essential for survival. In limbed animals, the frequency of locomotion is modulated primarily by changing the duration of the stance phase. The underlying neural mechanisms of this selective modulation remain an open question. Here, we report a neural circuit controlling a similarly selective adjustment of locomotion frequency in *Drosophila* larvae. *Drosophila* larvae crawl using peristaltic waves of muscle contractions. We find that larvae adjust the frequency of locomotion mostly by varying the time between consecutive contraction waves, reminiscent of limbed locomotion. A specific set of muscles, the lateral transverse (LT) muscles, co-contract in all segments during this phase, the duration of which sets the duration of the interwave phase. We identify two types of GABAergic interneurons in the LT neural network, premotor neuron A26f and its presynaptic partner A31c, which exhibit segmentally synchronized activity and control locomotor frequency by setting the amplitude and duration of LT muscle contractions. Altogether, our results reveal an inhibitory central circuit that sets the frequency of locomotion by controlling the duration of the period in between peristaltic waves. Further analysis of the descending inputs onto this circuit will help understand the higher control of this selective modulation.

## Introduction

Animals flexibly adapt their speed of locomotion to meet their behavioral needs (Alexander, 1989; Byrne, 2019; DeAngelis et al., 2019). In recent decades, the neural basis of the modulation of the speed of locomotion across the animal kingdom has received much attention. The mesencephalic locomotor region (MLR), which projects to reticulospinal neurons that in turn innervate spinal circuits, has been identified in all vertebrate species studied to date as an important control centre (Ryczko et al., 2017). Increasingly intense stimulation of the MLR causes increases in the speed of locomotion, with accompanying gait transitions (Atsuta et al., 1990; Grillner, 1985; Shik et al., 1966; Shik and Orlovsky, 1976; Skinner and Garcia-Rill, 1984). The spinal cord recruits different types of motor neurons at different speeds, with the accompanying changes in gait requiring widespread reconfiguration within its circuitry (Dasen, 2017; Kiehn, 2016).

How does the central nervous system vary the frequency of locomotion to achieve the required speeds? In a range of species, descending projecting excitatory neurons have been shown to drive the rhythm of locomotion (Berg et al., 2018; Caggiano et al., 2018; Capelli et al., 2017; Friesen and Kristan, 2007; Gatto and Goulding, 2018; Josset et al., 2018; Roberts et al., 2010). In mice, studies using optogenetics have shown that excitatory neurons are necessary and sufficient for rhythm generation (Hägglund et al., 2010, 2013), with studies ongoing to uncover the precise identity of the rhythm generators (Kiehn, 2016). Zebrafish, which use axial locomotion to move, have different central modules corresponding in adults to fast, intermediate, and slow locomotion that are selectively recruited to command the motor pools specific for different speeds (Ampatzis et al., 2013, 2014). The pacemaker neurons driving locomotion at different speeds have intrinsic bursting frequencies related to their module affiliation (Song et al., 2020).

The kinematics of movement change as a function of frequency depending on the species and gait. Swimming animals modulate their undulatory frequency by controlling the intersegmental lag, which is linearly scaled with the locomotor cycle duration (Grillner, 1974). On the other hand, limbed animals change the frequency of walking by varying the locomotor cycle differentially: the stance phase is varied, but the swing phase is almost unchanged, even as animals switch to different gaits. This holds true for animals ranging from insects and tardigrades to mammals (Boije and Kullander, 2018; Frigon et al., 2014; Grillner et al., 1979; Jacobson and Hollyday, 1982; Nirody et al., 2021). How the nervous system generates this asymmetry in the variation of stance and swing phases is still an open question (Bidaye et al., 2018; Boije and Kullander, 2018; Kiehn, 2016).

Here, we investigated the speed-dependent modulation of locomotion in *Drosophila* larvae and the underlying neural mechanisms. The *Drosophila* larva moves by peristaltic waves, in which body wall muscles contract sequentially from one end to the other (Berrigan and Pepin, 1995; Heckscher et al., 2012; Sun et al., 2022). We found that the *Drosophila* larval locomotor cycle is also differentially modulated: the phase in between each consecutive peristaltic wave (the “interwave” phase), not the peristaltic wave itself, is primarily varied with speed, reminiscent of the stance phase in limbed locomotion. We then examined the underlying muscular dynamics and found that the interwave phase is characterized by synchronous contractions of the lateral transverse (LT) muscles along the anterior-posterior axis. The amplitude and duration of their contraction scale with the duration of the interwave phase. Using EM connectomics and calcium imaging, we identified two types of interneurons that are associated with the LT neural circuitry and show segmentally synchronized activity: GABAergic premotor neuron A26f and its presynaptic partner GABAergic interneuron A31c. Using optogenetics, we revealed that both A31c and A26f neurons are sufficient and necessary for the desired contraction of the LT muscles and set the speed of locomotion through the modulation of the interwave phase. Our results reveal that the *Drosophila* larva uses a similar strategy to regulate speed as limbed animals by varying the two main phases of the cycle differentially and that the activity of an inhibitory circuit generates this variation.

## Results

### Variability in the interwave phase of crawling contributes to speed variability

Crawling behavior in *Drosophila* larvae is generated by repetitive waves of propagation along the length of their body (Berrigan and Pepin, 1995). A previous study in mildly physically restrained first-instar larvae showed that crawling speed correlates with stride period more than stride length (Heckscher et al., 2012). It has been shown that the lag between the contraction of adjacent segments during the peristaltic wave (intersegmental lag) scales with the cycle period in the intact animal and the isolated central nervous system (CNS) (Heckscher et al., 2012; Pulver et al., 2015). These observations suggest that the cycle period varies through a uniform, rather than an asymmetric, modulation of the phases of the locomotor cycle. However, these physically restricted larvae have long cycle periods (2-20 seconds), presumably due to aberrant or absent input from sensory neurons (Caldwell et al., 2003; Hughes and Thomas, 2007; Schützler et al., 2019; Zarin et al., 2019). How the larva varies locomotion in free crawling within the normal range of cycle periods (0.6-2 seconds) is therefore not understood. We therefore first aimed to confirm these findings in freely crawling third-instar larvae (Figure 1A-I, Figure 1-supplement A-F). We recorded larvae freely crawling on an agarose plate and measured the displacement of their body-wall segments (Figure 1A-C). Larvae crawled at varying speeds even within the same environmental conditions such as temperature (Figure 1D and 1E, 0.35 – 1.23 mm/sec, n = 18 larvae). We found that the locomotion speed in these freely crawling animals also correlated with stride frequency more so than stride length, consistent with the previous report (Figure 1E). To further characterize the underlying kinematic changes, we assessed how the two previously identified phases within the locomotor cycle (Heckscher et al., 2012) change with speed. In the first phase, local body wall contractions are propagated from the posterior to anterior segments (here called ‘wave phase’), whereas the second is characterized by the period from mouth parts unhooking to the onset of the tail contraction (‘interwave phase’; Figure 1F).

**Figure 1.**
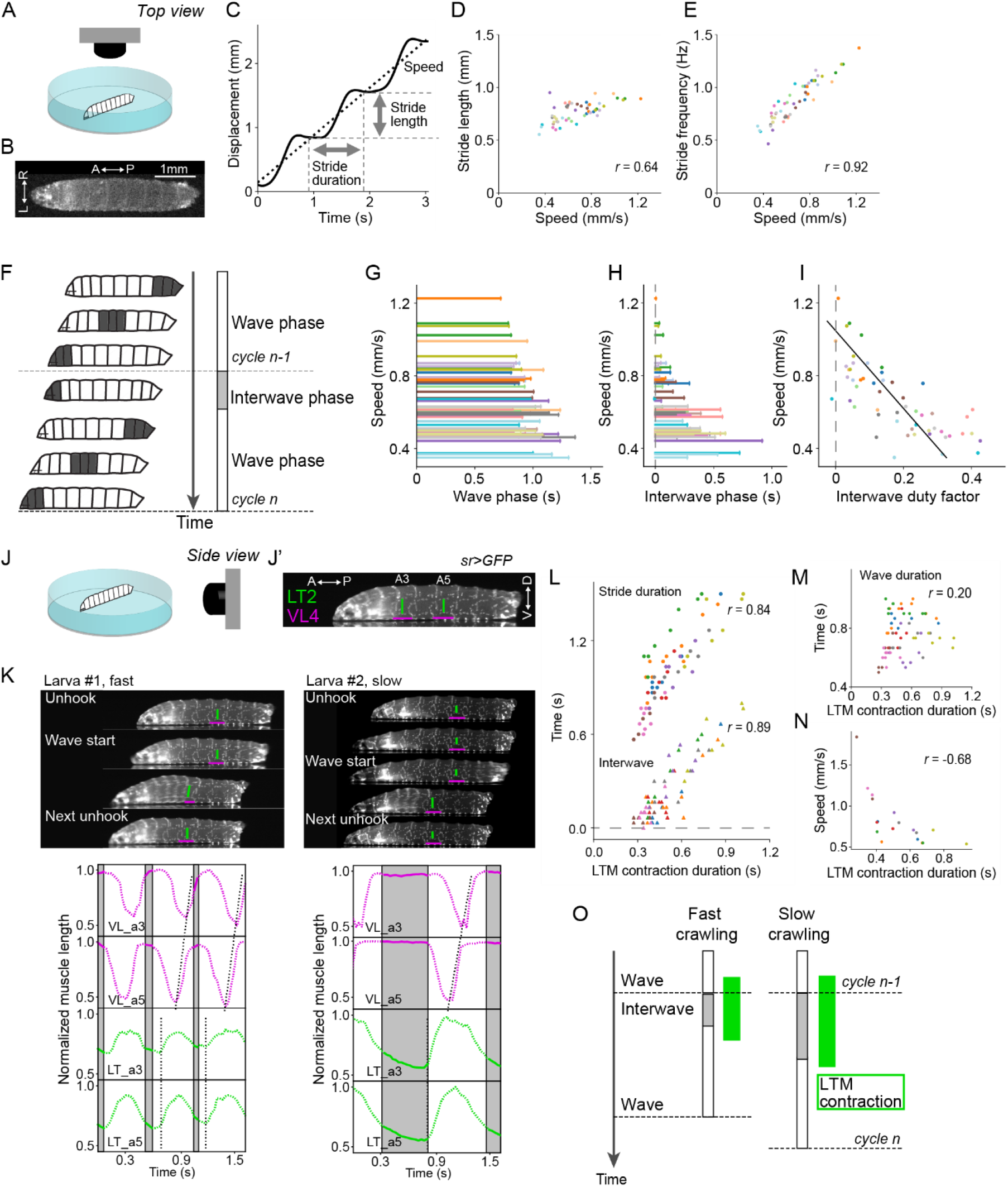
Crawling speed depends on the duration of interwave phase, during which the LT muscles are contracted. **(A-I)** Recording of locomotion parameters from top view (n = 268 strides, 54 episodes, 18 larvae). **(A)** Schematic drawing of the crawling assay from top view. **(B)** An example frame of top-view recording. **(C)** Measurement of the stride duration, stride length, and speed. **(D)** Relationship between speed and stride length. *r* represents Pearson correlation coefficient. **(E)** Relationship between speed and stride frequency. **(F)** Schematic representation of the two phases of a locomotor cycle. **(G)** Relationship between the duration of wave phase and speed. **(H)** Relationship between the duration of interwave phase and speed. **(I)** Interwave duty factor, the proportion of interwave phase in the stride duration, decreases with speeds. Linear regression coefficient is −0.47. **(J-O)** Recording of locomotion parameters and muscular kinematics from side view (n = 8 larvae). **(J)** Schematic drawing of the crawling assay from side view. **(J’)** An example frame of side-view recording. LT2: lateral transverse muscle 2. VL4: ventral longitudinal muscle 4. **(K)** Representative tracking of the muscle movement during forward crawling. Top left panel shows the muscle movement with a fast speed. Top right panel shows the muscle movement with a slow speed. Bottom panels demonstrate the dynamics of muscle lengths in the data shown in the top panels. **(L)** Relationship between the contraction duration of LT2 muscle and two temporal parameters (stride duration and interwave duration) (Pearson correlation coefficients; stride duration: 0.84 and interwave duration: 0.89). **(M)** Relationship between the contraction duration of LT2 muscle and wave duration (Pearson correlation coefficient: 0.20). **(N)** Relationship between LT2 muscle contraction duration and speed. **(O)** Schematic of the relationship between LT muscle (LTM) contraction and crawling speed. The duration of the two phases and the contraction of LT muscles are correlated with crawling speed.

To examine the possible contribution of the variability in the interwave phase to the speed variability, we analyzed the correlation between crawling speed and the two phases (Figure 1G-I). We found that both the interwave phase and the wave phase are correlated with the speed (Figure 1-supplement C and C’; Pearson correlation coefficient: the wave phase vs speed r = −0.62, the interwave phase vs speed r = −0.74). The interwave duration at faster speed is close to 0. Indeed, the duty factor of the interwave phase, which is given by the ratio of the interwave phase duration to stride duration, decreased with speed (Figure 1I; linear regression coefficient = −0.47, r^2^ = 0.55) and was reduced to zero at the faster speeds. The wave and interwave phases are modulated independently, as can be seen by the lack in correlation between these two phases (Figure 1-supplement D and E). These results suggest that the speed-dependent modulation of crawling frequency is largely due to modulation of the interwave phase.

We quantified the duration of each phase in freely crawling larvae. To evaluate the contribution of these two phases to stride duration, we plotted the duration of these phases as a function of stride duration (Figure 1-supplement F and F’). Both are correlated with stride duration, with the interwave phase correlated more strongly than the wave phase (interwave phase r = 0.86, wave phase r = 0.56, p < 0.0001). What becomes clear from this analysis is that when stride duration is less than approximately 1 second, the interwave phase is minimal, and the wave duration therefore reduces in duration in line with stride duration; on the other hand, when stride duration is greater than approximately 1.2 seconds, wave duration is more or less constant, with increases in stride duration accompanied by increases in interwave duration. These observations suggest that the interwave phase between peristaltic waves is more variable than wave phase, and that there is a range-dependent modulation of the frequency of locomotion.

### Synchronous contraction of transverse muscles is correlated with the intervwave phase

To reveal the nature of the interwave phase, we examined the movement of body wall muscles during free crawling. The ends of individual muscles were labelled by expressing GFP in the tendon cells using *sr-Gal4* (Schnorrer et al., 2007) and imaged from the side (Figure 1J and 1J’). This allowed us to analyze the contraction dynamics of each muscle in freely crawling larvae. The dynamics of two longitudinal muscles (DO1 and VL4) that span the anterior and posterior boundary of each segment and one transverse muscle (LT2) that runs perpendicular to the anterior-posterior axis of the animal were examined (Figure 1K and Figure 1-supplement G). Consistent with the previous study, longitudinal muscles exhibited propagation from the posterior segment to the anterior in forward crawling (Figure 1L and Figure 1-supplement G’). Interestingly, transverse muscles only showed synchronous contractions (Figure 1K and Figure 1-supplement G’). Furthermore, while longitudinal muscles were mostly contracting during peristaltic waves, the transverse muscles contracted during the interwave phase (Figure 1-supplement G’).

The phase-specific contraction of transverse muscles implies the possible involvement of transverse muscles during the interwave phase. Accordingly, we analyzed the contraction duration of transverse muscles and analyzed its relationship with the duration of the phases. While the duration of wave phase didn’t have a strong correlation, the duration of the interwave phase had a high correlation with the contraction duration of transverse muscles (wave phase r = 0.20, interwave phase r = 0.89, p < 0.0001, Figure 1L and 1M). Stride duration, which is the sum of the wave duration and the interwave duration, also had a strong correlation with the contraction duration of transverse muscles (r = 0.84, Figure 1L). This result implies that the contraction duration of transverse muscles should be related to the crawling speed. We therefore plotted the duration of transverse muscle contractions against crawling speed (Figure 1N) and found that they were correlated (r = −0.68). Next, we analyzed the relationship between the amplitude of transverse muscle contraction and the crawling kinematics. As is the case of the contraction duration, the contraction amplitude is also correlated with the interphase duration and stride duration but not the wave duration (Figure 1-supplement H-I). On the other hand, the contraction amplitude is weakly correlated with speed (Figure 1-supplement J). These results show that the duration and amplitude of synchronous contraction of transverse muscles are related to the duration of interwave phase (Figure 1O). Importantly, the duration of the synchronous contraction is correlated with crawling speed (Figure 1O).

### Identification of GABAergic interneurons A31c showing segmentally synchronized activity

In a screen for the neurons that are activated during the interwave phase, we identified cell type A31c (Figure 2-supplement). By reviewing the existing genetic driver expression patterns (Li et al., 2014), we identified several genetic drivers targeting the A31c neuron, including a split GAL4 driver (*A31c-a8-sp*) specifically targeting the A31c neuron in segment A8, a split GAL4 driver (*A31c-sp*) targeting A31c neurons in neuromeres A2-A8, and a LexA driver (*A31c-LexA*) that targets A31c neurons in neuromeres A2-A8 with variable expression patterns. We first used these lines to investigate the morphology and neurotransmitter identity of A31c neurons. The neurites project dorsally approximately one neuromere mostly to the anterior (Figure 2A). The synaptic input sites are located along the dorsolateral (DL) tract (Landgraf et al., 2003), while the synaptic output sites are mainly positioned dorsally near the midline (Figure 2A and 2B). Using immunohistochemistry, we found that A31c neurons are GABAergic (Figure 2-supplement A).

**Figure 2.**
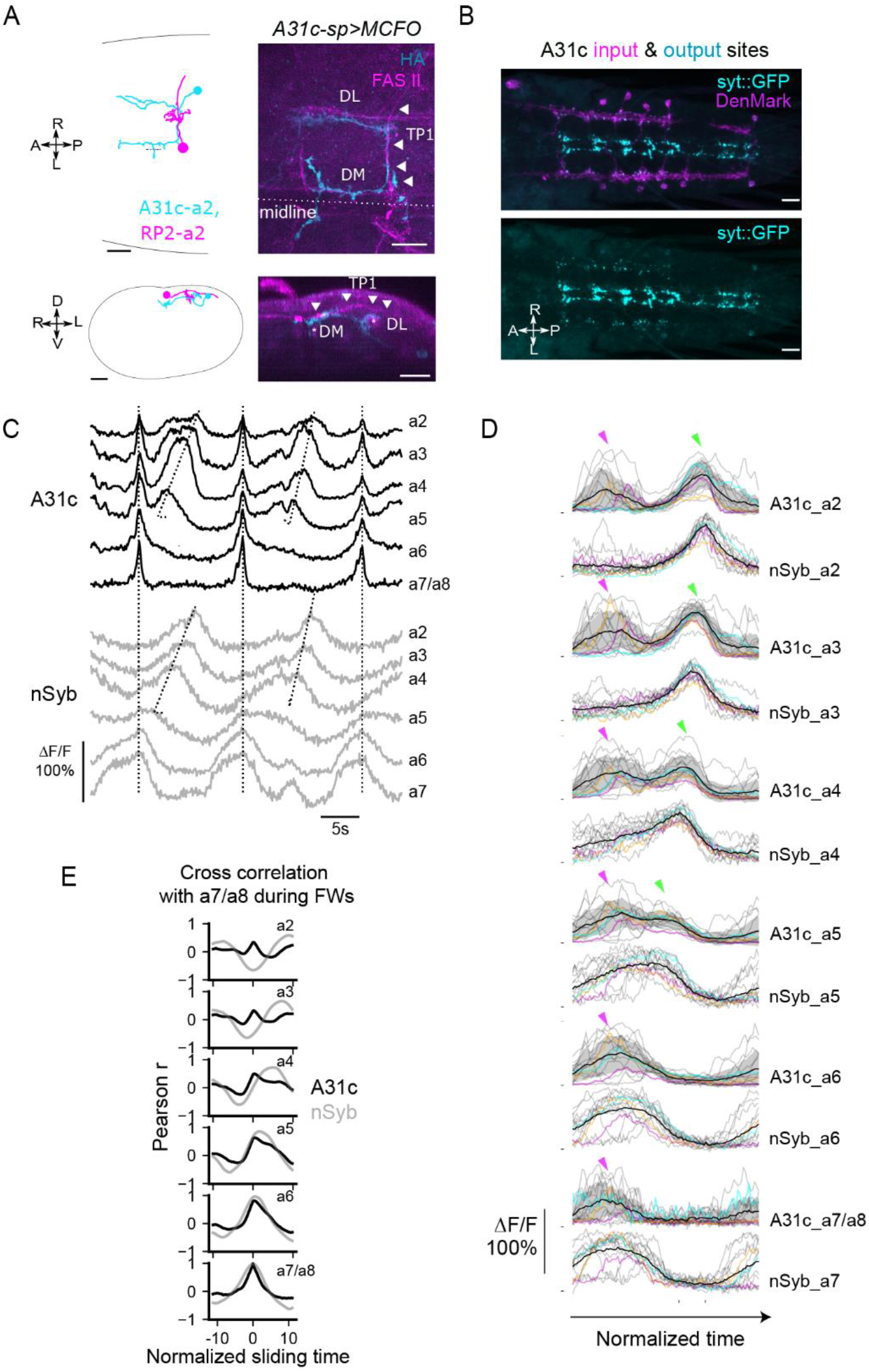
A31c neurons show synchronous activity preceding the forward wave. **(A)** A31c single-neuron morphology shown by EM reconstruction and genetically mosaic analysis. Scale bars: 20 μm. **(B)** Pre- and post-synapse markers label the input and output sites of A31c neurons (*A31c-sp>UAS-syt::GFP, UAS-DenMark*). Scale bars: 20 μm. **(C-E)** Recording of calcium activity of A31c neurons (*A31c-sp>UAS-CD4::GCaMP6f*) and group activity of nSyb neurons (*nSyb-LexA>LexAop-RGECO1*) which reports the pan-neuronal activity in fictive locomotion (n = 15 traces, 5 larvae). **(C)** Example recordings of A31c neurons and nSyb neurons in fictive forward locomotion. **(D)** Group data of calcium imaging of A31c neurons and nSyb neurons. Each trace is aligned by activity peak of nSyb_a4 and nSyb_a2 and normalized to 0-1 by the activity maximum and minimum of the whole recording. Magenta arrows indicate the co-activation of A31c neurons. Green arrows indicate the wave-like activity of A31c neurons. Black lines represent the average calcium activity. Shading represents the standard error. Colored lines represent the three example traces. Grey lines represent all other traces. Ticks along the horizontal axis indicate the activity peaks of nSyb_a4 and nSyb_a2. Ticks along the vertical axis indicate the 0. **(E)** Cross correlation of neuronal activity between the neuron in each segment (from A2 to A7/A8) and the one in A7/A8 (black: A31c neurons, grey: nSyb neurons). See Materials and methods for details.

We then used a dual-color imaging system to monitor the activity of A31c neurons using *A31c-sp>UAS-CD4::GCAMP6f* and the pan-neuronal activity using *nSyb-LexA*>*LexAop-RGECO1* in the isolated CNS (Figure 2C-E and Figure 2-supplement B-C; Materials and methods for details). The pan-neuronal activity patterns were used as an indicator of the fictive behaviors produced (Pulver et al., 2015), showing stereotyped fictive forward waves (FW). At the initiation of forward locomotion, all abdominal A31c neurons show burst-like coactivation preceding the forward wave (Figure 2C and 2D). During the FW that follows, A31c neurons in anterior segments A2-A5 are re-activated in a wave-like sequence (Figure 2C and 2D). The intersegmental lags of pan-neuronal activity between neighboring segments show non-zero values which reflects the propagation of neuronal activity along the body axis (Figure 2E). On the other hand, the intersegmental lags of A31c neurons are almost zero consistent with their synchronized activity (Figure 2E). To sum, these results show that A31c neurons exhibit synchronous multi-segmental activity during the interwave phase.

### A31c neurons receive synaptic inputs from descending neurons and give synaptic output to local and ascending neurons

To understand the details of the connectivity of this circuit, we reconstructed the connectivity of A31c using EM connectomics (Materials and methods for details). We identified A31c neurons in neuromeres A2-A8 in the database of the larval central nervous system, reconstructed all pre- and post-synaptic partners, and analyzed their connectivity (Figure 3A-3C and Figure 3-supplement A and B). We analyzed the connectivity of A31c neurons in anterior segments A2-A3 and posterior segments A7-A8 separately (Figure 3B-3D). We found that the synaptic inputs to A31c neurons are similar in the anterior and posterior segments, with several descending cell types innervating A31c across segments (Figure 3D). The same suboesophageal (SEG) descending neuron cell type (here labelled “S10”) provides a significant plurality of the synaptic input. Among postsynaptic targets, we found that just one cell type is consistent between segments A2-A3 and A7-A8: A26f neurons, which are among their top three postsynaptic partners (Figure 3D). A26f neurons also receive synaptic inputs from the ascending cell type A19f, one of the top postsynaptic partners of A31c neurons (Jonaitis, 2020). Interestingly, it has previously been reported that A26f strongly innervates the transverse motor neurons (Zarin et al., 2019; Zwart et al., 2016).

**Figure 3.**
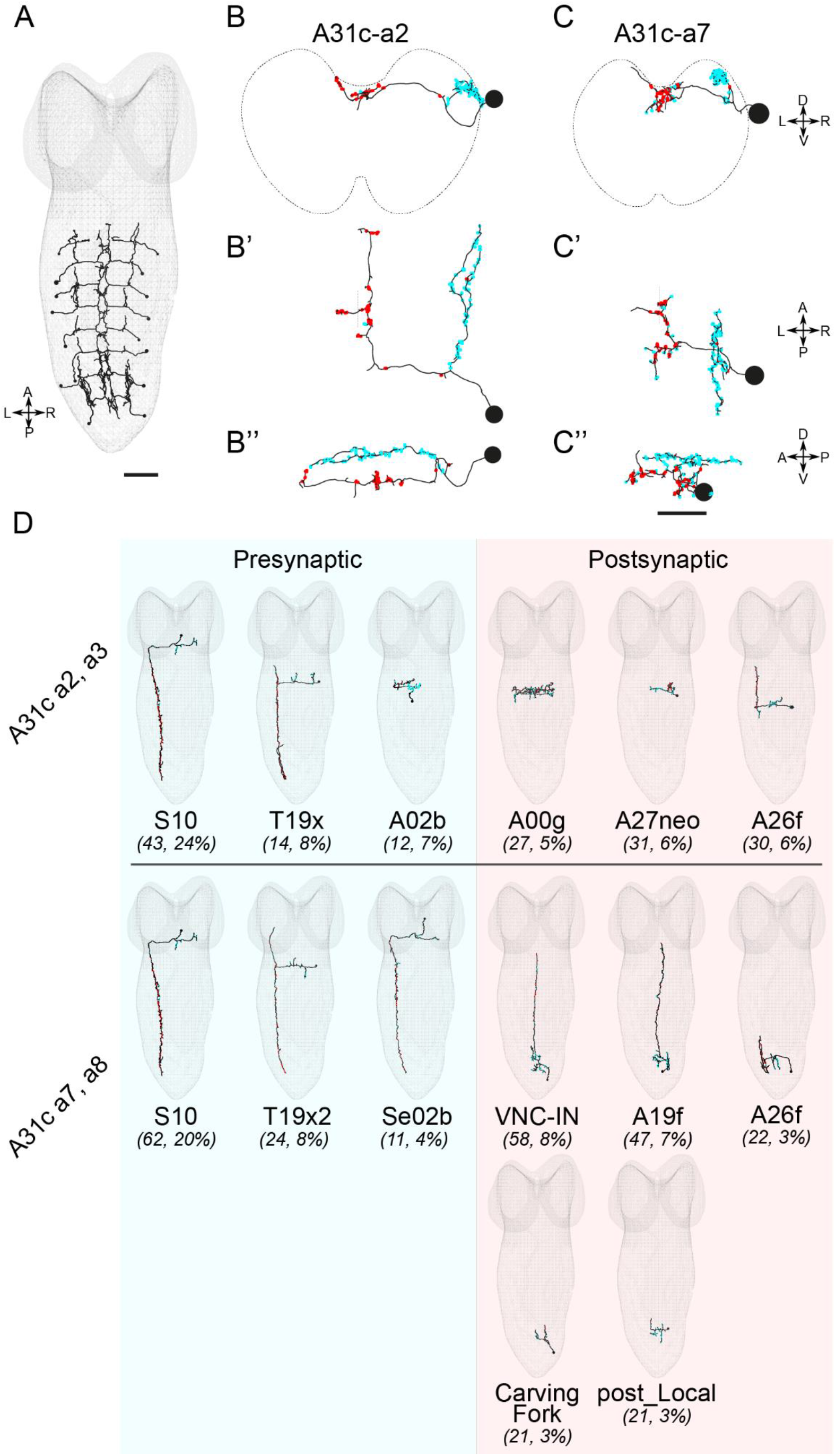
EM reconstruction reveals the connectivity of A31c neurons. **(A)** Identification of all A31c neurons in the EM database. Scale bars: 10 μm. **(B)** Morphology of an anterior and a posterior A31c neuron. Red indicates the output sites. Cyan indicates the input sites. **(C)** Example EM images of a presynaptic site of A31c neuron. Scale bar: 0.5 μm. **(D)** Top pre- and post-synaptic partners of anterior and posterior A31c neurons.

We next used *trans*-Tango, a genetic tool for tracing postsynaptic partners (Talay et al., 2017), to confirm the identity of the postsynaptic neurons of A31c-a8. We repeatedly identified Tango expression in an A26f-like cell type in segment A7, in addition to other neurons, some of which we could identify (four samples showing A26f-a7 neurons; Figure 3-supplement C and D). These results collectively show that A26f neurons are postsynaptic to A31c neurons.

### Identification of GABAergic premotor neurons A26f showing segmentally synchronized activity preceding the fictive forward wave

Since A26f neurons are postsynaptic to A31c (Figure 3D and Figure 3-supplement C and D) and strongly innervate LT motor neurons (Figure 4A and Figure 4-supplement A; Zarin et al., 2019, Zwart et al., 2016), we next focused on A26f neurons to understand the neural mechanism underlying the generation of the interwave phase. We used a split GAL4 driver (“*A26f-sp*”), which labels A26f neurons in neuromeres A3-A5 (Figure 4B), to investigate their morphology and neurotransmitter identity. A26f neurons have synaptic input sites dorsally near the midline and synaptic output sites near the DL tract (Figure 4B). Remarkably, A26f neurons project their axon along the DL tract for multiple segments. As a representative example, the axon of A26f neuron of neuromere A5 extends four neuromeres from A6 neuromere to A3 neuromere (Figure 4B). We found that A26f neurons are GABAergic (Figure 4-supplement B). It has previously been reported that A26f neurons are corazoninergic (Zarin et al., 2019). However, a comparison of the morphology of A26f neurons with confirmed corazoninergic neurons (Santos et al., 2007) and the absence of peptidergic dense core vesicles in A26f neurons in the EM connectomics dataset suggest that A26f neurons are not corazoninergic. A26f neurons form inhibitory synapses to MNs innervating LT muscles in multiple neuromeres, which suggests the potential of the A26f neurons to control the activity of LT muscles broadly in multiple segments.

**Figure 4.**
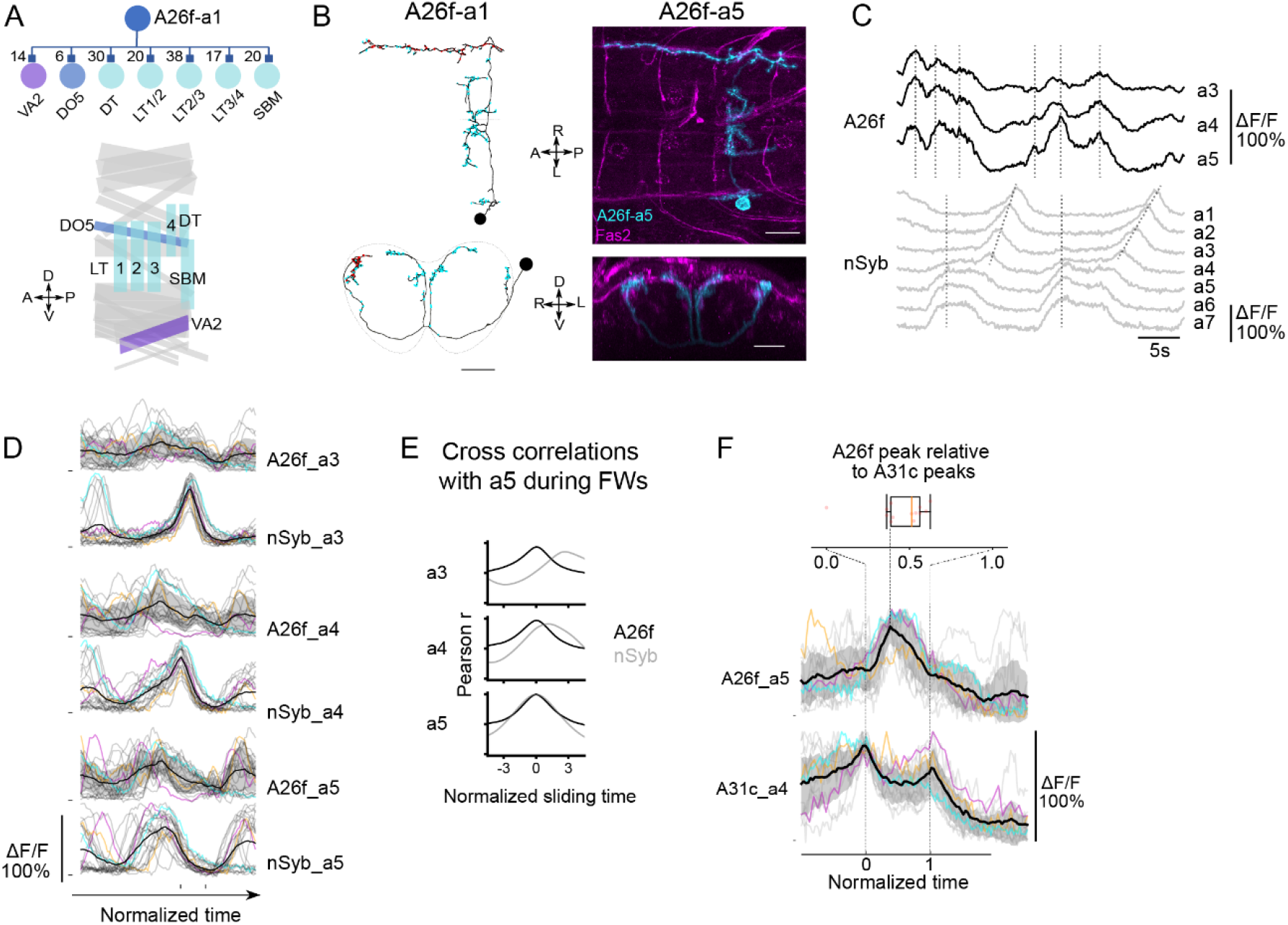
A26f neurons inhibit motor neurons and exhibit synchronous activity at the initiation of forward locomotion. **(A)** A26f neurons innervate motor neurons. (Top) Postsynaptic neurons of A26f neurons revealed by connectomics analysis in A1 neuromere (Zarin et al., 2019). (Bottom) Layout of body wall muscles in a hemi-segment. Purple, blue, and sky blue muscles are innervated by motor neurons in the same color in the top panel. **(B)** Morphology of A26f neurons shown by the EM reconstruction and confocal images. Scale bars: 20 μm. **(C)** Recording of calcium activity of A26f neurons (*A26f-sp>UAS-CD4::GCaMP6f*) and group activity (*nSyb-LexA>LexAop-RGECO1*). **(D)** Group data of calcium imaging of A26f neurons and nSyb neurons. Each trace is aligned activity peak of nSyb_a4 and nSyb_a2 and normalized to 0-1 by the activity maximum and minimum of the whole recording. Black lines represent the average calcium activity. Shading represents the standard error. Colored lines represent the three example traces. Grey lines represent all other traces. Ticks along the horizontal axis indicate the activity peaks of nSyb_a4 and nSyb_a2. Ticks along the vertical axis indicate the 0. **(E)** Cross correlation of neuronal activity between the neuron in each segment (A3-A5) and the one in A5 (black: A26f neurons, grey: nSyb neurons). See Materials and methods for details. **(F)** Simultaneous calcium imaging of A31c and A26f neurons. (Bottom) Recording of calcium activity of A26f-a5 neuron (*A26f-sp>UAS-CD4::GCaMP6f*) and its presynaptic partner A31c-a4 neuron (*A31c-LexA>LexAop-jRCaMP1b*) (n = 13 traces, 7 larvae). Colored lines indicate the example traces. Black lines indicate the average calcium activity. Grey lines indicate all other traces. (Top) Peak time of A26f signals relative to the first peak time of A31c signals.

Next, we related the activity patterns of A26f neurons to behavior by performing dual-color imaging experiments of *A26f-sp*>*UAS-CD4::GCaMP6f* and *nSyb-LexA*>*LexAop-RGECO1* in the isolated CNS (Figure 4C and Figure 4-supplement C and D; Materials and methods for further details). Unlike most neurons showing fictive wave-like activity (Lemon et al., 2015), A26f neurons only have synchronized activity in neuromeres A3-A5, which mostly occurred during the periods out of the fictive waves (Figure 4C and 4D). Consistent with synchronized activity, there are high correlations between the activity of A26f segmental homologs, unlike pan-neuronal activity (Figure 4E). The A26f neurons can exhibit one or several peaks at the initiation phase of the FW (Figure 4C and 4D). To sum, A26f neurons have four important characteristics: (1) They form inhibitory synapses with motor neurons in multiple segments targeting transverse muscles; (2) A26f neurons in the abdominal segments are activated simultaneously; (3) A26f neurons are activated between wave phases; (4) they are postsynaptic to A31c neurons.

As both A26f and A31c neurons show robust synchronous activity at the initiation of FW, we then monitored the activity of the two neurons simultaneously by using *A31c-LexA>LexAop-jRGECO1b, A26f-sp>UAS-CD4::GCaMP6f*. We found that the synchronous peak of A26f neurons is “bookended” by the peaks in A31c activity of neighboring segments (Figure 4F). This is consistent with the inhibitory nature of the A31c-A26f synapses and suggests these cell types might be involved in determining the duration of the interwave phase.

### Activation of A26f neurons reduced the amplitude of LT muscle contraction during the forward crawling

To assess whether A26f neurons can inhibit the activity of LT muscles, we analyzed muscle responses to the optogenetic activation of A26f neurons in forward cycles. We combined the optogenetic activator *UAS-CsChrimson* targeted by *A26f-sp* to activate the A26f neurons and the muscle genetic marker *mhc-GFP* expressing GFP to visualize the body wall muscles (*A26f-sp>UAS-CsChrimson, mhc-GFP*). We used *A26f-sp* negative animals as a control (*UAS-CsChrimson, mhc-GFP*). Because of the spectral overlap between the light to activate CsChrimson and that to excite GFP, we used a confocal microscopy system that separates the light for optogenetics and imaging into two sections of the objective back aperture, respectively in combination with a new preparation called sideways preparation (Figure 5A and 5B; Materials and methods for detail).

**Figure 5.**
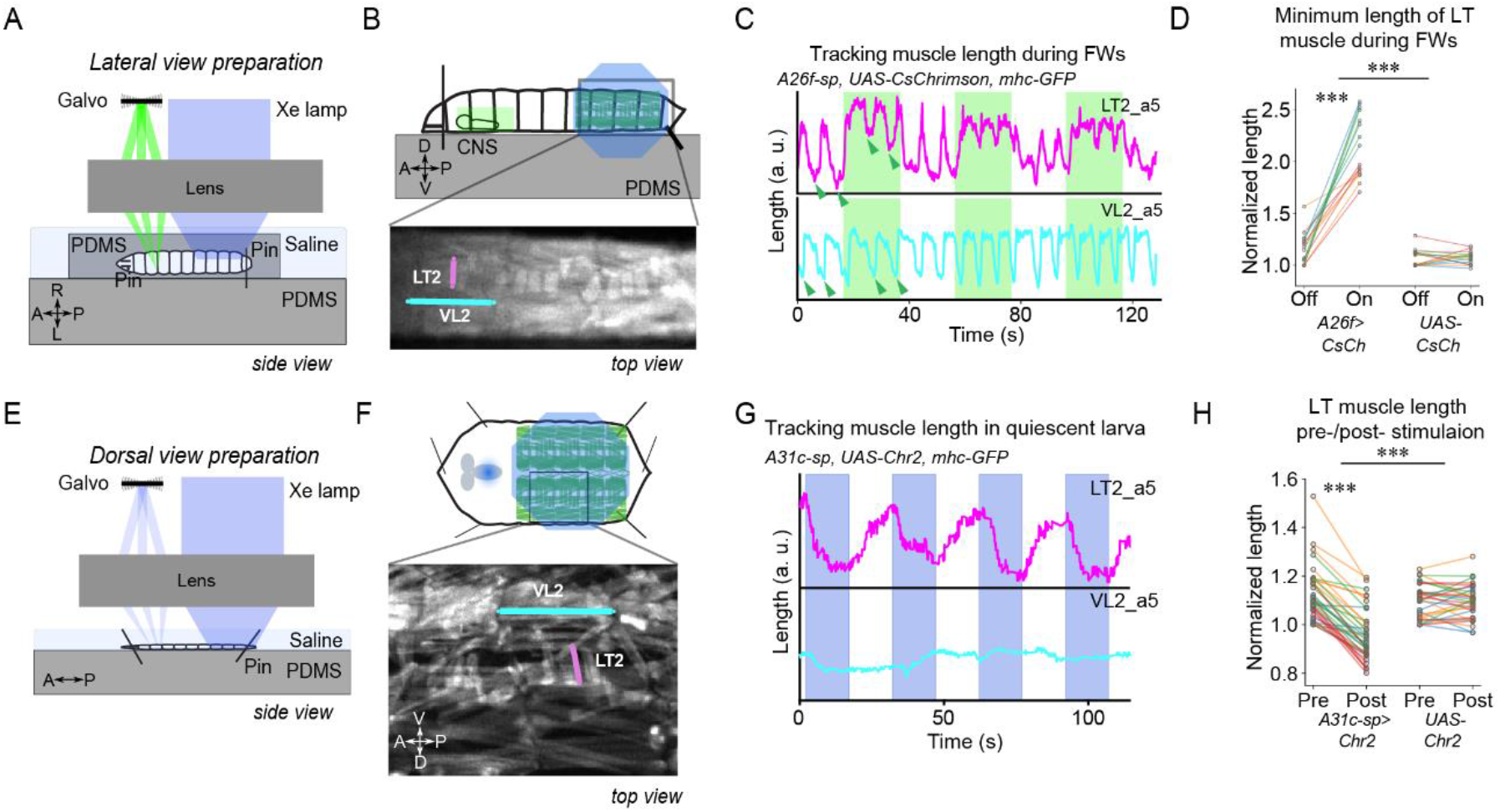
Optogenetic activation of A31c or A26f neurons affects the contraction of LT muscles. **(A-D)** Optogenetic activation of A26f neurons reduces the contraction amplitude of the LT2 muscle during forward crawlings in the sideways preparation. **(E-H)** Optogenetic activation of A31c neurons causes contraction of the LT muscle in the fillet preparation. **(A and E)** Experimental setups. See Materials and methods for details. **(B and F)** Schematics of the imaging setup (top) and sample fluorescence images (bottom). **(C and G)** Traces of the length of the transverse muscle LT2 and the longitudinal muscle VL2 in the optogenetic experiments. Shaded regions show the timing when the light stimulus is applied. Arrowheads indicate where the measurement was made in Figure 5D. **(D)** The minimum length of the LT2 muscle was increased by the activation of A26f neurons. Muscle lengths are normalized to the minimum length during the light-off period. The hierarchical bootstrap test (See Material and methods for details.) **(H)** The length of the LT2 muscle in the resting state was decreased by the activation of A31c neurons. Muscle lengths are normalized to the minimum length during the light-off period. The hierarchical bootstrap test (See Material and methods for details.)

We tracked the length of muscle LT2 and longitudinal muscle VL2 in segment A5 upon optogenetic stimulation. Activation of the A26f neurons reduced the contraction amplitude of the LT2 muscle, while the contraction of the VL2 muscle was almost unchanged (Figure 5C, 5D, and Figure 5-supplement A). These results confirm that activation of A26f neurons is sufficient for the inhibition of the contraction of LT muscles.

### Activation of A31c neurons induced the contraction of LT muscles

As A31c could inhibit A26f based on the connectivity (Figure 3D and Figure 2-supplement A) and activity (Figure 4G’) analyses described above, we tested whether activation of A31c neurons can enhance contractions of LT muscles. We therefore activated A31c neurons and analyzed the change in LT2 muscle length using animals carrying *A31c-sp>UAS-Chr2*.*T159C, mhc-GFP* transgenes. We restricted the stimulation laser to the abdominal neuromeres in a semi-intact preparation (“fillet preparation”, Pulver et al., 2015) to avoid activating SEG or brain neurons (Figure 5E and 5F). The stimulation caused the contraction of LT muscles in all visualized abdominal neuromeres (A3-A8; Figure 5G), causing a reduction in the minimum length of the LT2 muscle (Figure 5H). These results suggest that activation of A31c neurons is sufficient to activate the LT muscles. As no apparent contraction of other muscles was observed (Figure 5-supplement B), we assume that the A31c neurons mainly regulate the activity of the LT muscles.

### Silencing A31c or A26f neurons influenced the amplitude of the LT muscle contraction during forward crawling

Next, we examined if the interneurons of interest were required for the observed contraction of the LT muscles. To test this, we used optogenetic silencing combined with muscular imaging in the sideways preparation (Figure 5A). We first tested whether A26f neurons are required for the contractions of transverse muscles by using animals carrying *A26f-sp>UAS-GtACR1, mhc-GFP* for optogenetic silencing and muscular visualization. We found that the minimum length of muscle LT2 decreased after optogenetic silencing of A26f, suggesting increased levels of contraction (Figure 6A and 6B). We next assessed the requirement of A31c neurons by using *A31c-sp>UAS-GtACR1, mhc-GFP*. We found that after optogenetic silencing, the minimum length of the LT2 muscle in segment A5 was increased during forward cycles (Figure 6A and 6B). The minimum length of transverse muscle VL2 was not affected by the inhibition of A26f or A31c (Figure 6C). These results reveal that the activity of A26f and A31c neurons is both necessary and sufficient for the appropriate contractions of LT muscles observed during locomotor cycles.

**Figure 6.**
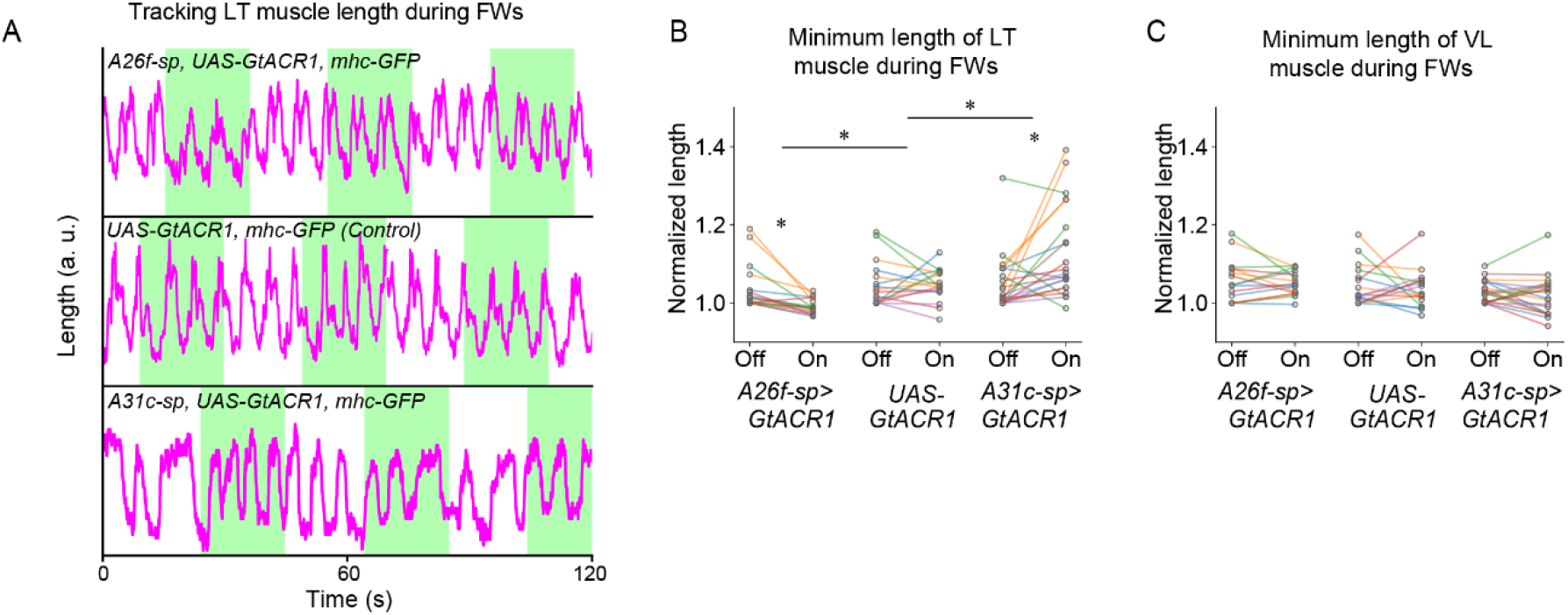
Optogenetic inhibition of A31c or A26f neurons affects the contraction of LT muscles. **(A)** Traces of the length of transverse muscles (LT2) in the sideways preparation with optogenetic stimulation. **(B)** The minimum length of LT muscles was affected by the optogenetic inhibition of A31c or A26f neurons. Muscle lengths are normalized to the minimum length during the light-off period. The hierarchical bootstrap test (see Material and methods for details.) **(C)** The minimum length of VL muscles was not affected by the optogenetic inhibition of A31c or A26f neurons. Muscle lengths are normalized to the minimum length during the light-off period.

### A26f neurons modulate interwave duration

Our previous results suggest that the activation of A26f neurons reduces the contraction of the LT muscles, thereby potentially reducing the duration of the interwave phase. To test this hypothesis, we activated A26f neurons and analyzed the kinematics of crawling in animals of the genotype *A26f-sp>CsChrimson* on low-concentration agarose plates (0.7%) (Figure 7A and 7B). We used animals that lacked the *A26f*.*DBD* transgene as a control (*A26f*.*AD>CsChrimson*).

**Figure 7.**
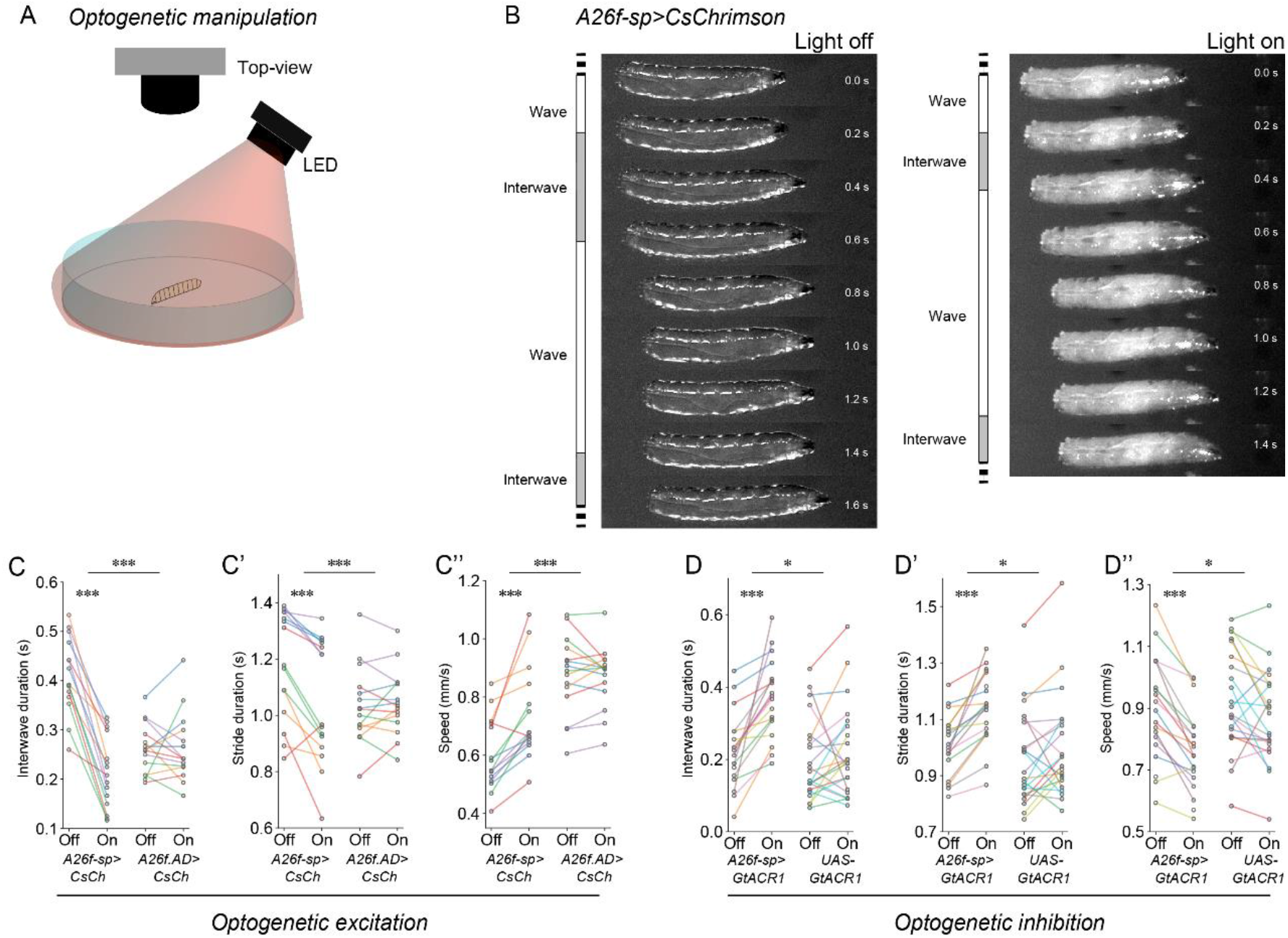
Optogenetic manipulation of A26f neurons affects the interwave duration. **(A)** Experimental setup for optogenetics in free-crawling larvae. **(B)** Example frames show that the interwave phase is reduced during the activation of A26f neurons. **(C-C’’)** Crawling dynamics changed during the activation of A26f neurons. The hierarchical bootstrap test (See Material and methods for details.) **(D-D’’)** Crawling dynamics changed during the inhibition of A26f neurons. The hierarchical bootstrap test (See Material and methods for details.)

During optogenetic activation of A26f neurons, larvae exhibited faster crawling (Figure 7B). By analyzing the kinematics, we confirmed that the interwave phase and the total stride duration were both significantly decreased during the optogenetic activation (Figure 7C and 7C’). On the other hand, the wave duration was slightly increased (Figure 7-supplement A). Consistent with these results, the speed of crawling was significantly increased (Figure 7C’’). These results suggest that the activation of A26f neurons is sufficient to increase stride frequency and speed.

Next, we asked if A26f neurons are required to regulate the interwave phase and thereby the speed of freely crawling animals. To this end, we optogenetically inhibited A26f neurons in animals carrying *A26f-sp>GtACR1* and analyzed their crawling kinematics (Figure 7D-D’’ and Figure 7-supplement C and D). We found that inhibiting the A26f neurons increased the interwave duration and stride duration but had no significant effect on the wave duration (Figure 7D, 7D’ and Figure 7-supplement C), resulting in decreased speeds (Figure 7D’’). Combined with our previous analyses, these results indicate that the A26f neurons are functionally required to regulate the speed of locomotion by modulating the contraction of LT muscles.

### Activation of A31c neurons caused the increase in the interwave duration and the stride duration

Next, we tested the effect of manipulating A31c on crawling (Figure 7-supplement C-L). To activate A31c neurons, we used a genetic system *UAS-VNC-CsChrimson* that confines the expression of CsChrimson to the VNC neurons targeted by the *A31c-sp* transgene, resulting in expression in neuromeres A2-A8. We used animals carrying *UAS-VNC-CsChrimson* as a control. During the activation of A31c neurons, the interwave duration and the stride duration were significantly increased, while no significant difference was found in the wave duration (Figure 7-supplement E, F, and H). These effects are consistent with those observed in the inhibition of A26f neurons (Figure 7D, 7D’, and Figure 7-supplement C). On the other hand, the perturbation of A31c neurons could not induce other phenotypes in crawling kinematics (Figure 7-supplement G and I-L), which implies the involvement of other presynaptic neurons to A26f neurons in speed control. Consequently, these data imply that A31c neurons should contribute to the regulation of interwave phase duration through A26f neurons by the multi-segmental synchronous excitation within the interwave phase.

## Discussion

In summary, we aimed to understand the neural mechanisms that underlie the selective modulation of one phase of the locomotor cycle. We used the *Drosophila* larva as a model and found that this animal uses a strategy to primarily vary the phase between consecutive peristaltic waves for speed regulation. To implement this strategy, the larva modulates the amplitude and duration of the contraction of the LT muscles that are perpendicular to the crawling direction, which contract synchronously along the anterior-posterior axis before the onset of the peristaltic wave. The GABAergic interneurons A26f and A31c, upstream of the MNs innervating the LT muscles, showed segmentally synchronized activity preceding the FW. Connectivity analysis further revealed that A31c neurons receive shared descending input, and synapse onto ascending neurons and local neurons, including A26f. A26f neurons are sufficient and required for the desired activity of the LT muscles and thus the interwave duration and speed. Altogether, we established a neural basis for speed regulation by linking the speed-dependent modulation of contractions of muscles to the interneurons that control their activity (Figure 8).

**Figure 8.**
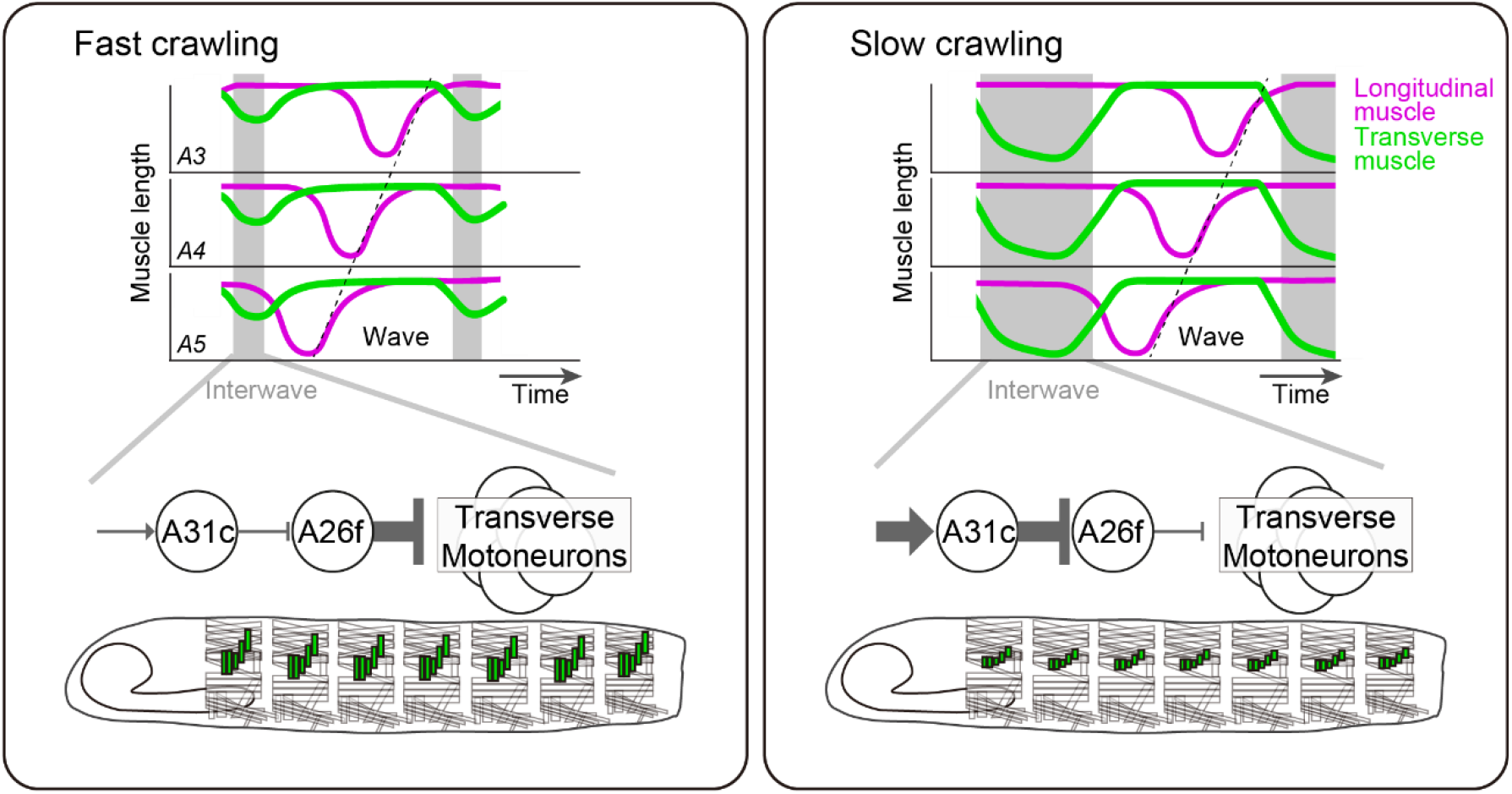
Schematics of larval speed control. Contraction of transverse muscles is suppressed by A26f neurons to make the interwave phase short and crawl fast (left panel). Multi-segmental synchronous activity of A31c and A26f neurons elongates the interwave phase duration to decrease crawling speed (right panel).

### Mechanics/mechanisms of modulating speed of locomotion

Locomotion speed is a function of the frequency and length of a cycle. We found that, similar to previous findings, the speed of larval crawling is determined more so by stride frequency than stride length (Frigon et al., 2014; Grillner et al., 1979; Jacobson and Hollyday, 1982; Nirody et al., 2021). Furthermore, similar to limbed animals including mammals and other insects, the two constituent phases of a locomotor cycle vary differentially with speed, with the interwave phase varying more than the wave phase. This similarity between the dynamics of locomotion between limbed and axial locomotion could be indicative of the kinematic constraints of each type of movement. For limbed locomotion, the forces required to move the limb during the swing phase depend on the limb’s mass. Large limbed animals, such as horses and humans (Boije and Kullander, 2018; Minassian et al., 2017), could use momentum-based strategies, requiring only brief active contractions in swing muscles, whereas smaller animals such as stick insects and mice (Bellardita and Kiehn, 2015; Bidaye et al., 2018) require constant neural input onto the swing muscles. In both cases, the relative invariance of the duration of this phase suggests that the rotational inertia of swinging limbs may be an important limiting factor in limbed locomotion (Kilbourne, 2013; Kilbourne and Hoffman, 2013, 2015; Rocha-Barbosa et al., 2005). The relative invariance of the wave phase suggests a similar constraint on the *Drosophila* larval motor system. Furthermore, the transverse muscle contractions during the interwave phase may be an evolutionary adaptation, with an additional set of neurons that control it. Consistent with this hypothesis, the last common ancestor to all bilaterians, the so-called “Urbilaterian”, is thought to have only had circular and longitudinal muscles (Cannon et al., 2016), and transverse muscles have only been identified in the larvae of some other species of Diptera (e.g., *T. castaneum*, Schultheis et al., 2019; *G. mellonella*, Emery et al., 2019). Furthermore, the transverse muscles are under independent neuromodulatory control (Elliott et al., 2021), and their motor neurons are innervated by a distinct set of interneurons (Zwart et al., 2016, Kohsaka et al., 2019).

In addition to the kinematic and evolutionary constraints, the differential modulation of the locomotor cycle may have several advantages. First, it could be a more efficient control strategy: independent control of the extension of the head and the peristaltic wave may reduce the complexity of motor control, and increase the flexibility of the head and tail by allowing the two ends to be moved separately. This may be particularly important as the anterior-most segments are involved in other motor programs including feeding (Melcher and Pankratz, 2005), and the transverse muscles control self-righting behaviour (Picao-Osorio et al., 2015). Second, it could improve energy efficiency within a particular range of speeds: though the contraction and extension of LT muscles during the movement of the head must entail a metabolic cost, the energy cost of moving the center of mass (CoM) might be reduced during slower movements. The CoM is mainly moved in the pistoning phase during the head extension and the tail contraction (Heckscher et al., 2012), and the transverse muscle contractions, which we speculate are involved in driving head movements, might be a more efficient method of extending the anterior segments by regulating the hydrostatic skeleton (Trimmer and Issberner, 2007).

### Neuronal control of speed modulation

A large body of work has identified the neural basis of the regulation of speed of locomotion in vertebrates, identifying associated circuits across the brain and spinal cord. Mouse, zebrafish, and Xenopus spinal cord preparations have been used to describe the selective recruitment of specific interneuron and motor neurons at different speeds (Berg et al., 2018; Boije and Kullander, 2018; Gatto and Goulding, 2018; Grillner and El Manira, 2020; Grillner and Kozlov, 2021; Kiehn, 2016; Roberts et al., 2010). These neurons are interconnected between members of the same “module”, each of which is sequentially recruited as the animal adjusts its speed of locomotion. Neuromodulation tunes the recruitment of neurons within these modules in the adjustment of speed during locomotion (Jha and Thirumalai, 2020). In limbed animals, changes in speeds are often accompanied by changes in gait (Bellardita and Kiehn, 2015). How the corresponding qualitative and quantitative changes in the locomotor cycle are achieved, is an area of active research. In the mouse spinal cord, V2a interneurons are required for maintaining left–right alternation at high-speed trotting (Crone et al., 2009). Furthermore, commissural V0V neurons are necessary for trot at all speeds, and ablation of commissural V0V and V0D neurons abolishes walk, trot, and gallop gaits (Bellardita and Kiehn, 2015). In the brainstem, the mesencephalic locomotor region (MLR) controls the initiation of locomotion and the expression of specific gaits. The MLR’s cuneiform nucleus (CnF) and the pedunculopontine nucleus (PPN) mediate alternating locomotor stepping in mice, whereas the CnF alone is necessary for high-speed synchronous locomotion such as found in galloping (Caggiano et al., 2018). A recent study identified distinct subclasses of glutamatergic neurons within the MLR, each with distinct roles in motor control outside of locomotion (Ferreira-Pinto et al., 2021), suggesting this nucleus has wider roles in regulating behavior. Despite this recent attention to the modulation of the speed of locomotion, the neural basis of the differential modulation of the locomotor cycle is still unknown. We have uncovered a set of inhibitory neurons, whose activity determines the duration of the interwave phase, thereby setting the frequency of locomotion. The inhibitory nature of this set of cells to regulate muscle contractions has many parallels in other systems. One of the simplest circuit designs for rhythm generation, the ‘half-center oscillator’, relies on reciprocal inhibition to generate alternating patterns of activity (Marder and Bucher, 2001), and reciprocal inhibition within the spinal cord is thought to underlie the generation of alternation during locomotion (Deliagina and Orlovsky, 1980; Geertsen et al., 2011; Pratt and Jordan, 1987). Indeed, inhibitory neurons shape the rhythms of neural activity on different timescales in systems from crustacean stomatogastric ganglion to vertebrate cortical circuits (Cardin, 2019; Marder and Bucher, 2001). In addition, a parallel between the *Drosophila* larval system and limbed locomotion can be seen in the mechanics of movement. For instance, in cats, the extensor muscles are mainly activated during the stance phase, while the flexor muscles are mainly activated in the swing phase (Engberg and Lundberg, 1969); similarly, we found that the fruit fly larva contracts its transverse muscles during the interwave phase, and its longitudinal muscles during the wave phase. These parallels may be mirrored within the neural circuitry mediating these muscle contractions. While the detailed implementation will obviously differ, the inhibitory neural circuit motif underlying the generation of the asymmetry of the two constituent phases of locomotion could therefore be conserved between species.

## Materials and methods

### Fly strains

Except where specifically mentioned, larvae were raised in standard cornmeal-based food at room temperature (25 °C), and third instar larvae were used for experiments. We used the following *all-trans* retinal (ATR) feeding conditions for optogenetics: 10 mM ATR yeast from 18 to 36 hours in CsChrimson and Channelrhodopsin 2 (Chr2.T159C) groups, 3 mM ATR yeast from 24 to 48 hours in GtACR1 groups. Fly strains are listed in Table 1. We used the split GAL4 drivers *A31c-a8-sp* (*R24H08-GAL4*.*AD, R45F08-GAL4*.*DBD*), *A31c-sp* (*R41F02-GAL4*.*AD, R44F09-GAL4*.*DBD*), and *A26f-sp* (*VT050223-GAL4*.*AD, R15E05-GAL4*.*DBD*). Transgenic flies *nSyb-LexA* were generated in the lab. The enhancer sequence of *neuronal Synaptobrevin (nSyb)* (*R57C10*, Pfeiffer et al., 2012) was cloned into pBPLexA::p65Uw plasmid (Pfeiffer et al., 2010). The transgenic line was generated in the *VK00027* locus (BestGene Inc., USA). Sources of the fly strains are listed in Table 1.

**Table 1.**
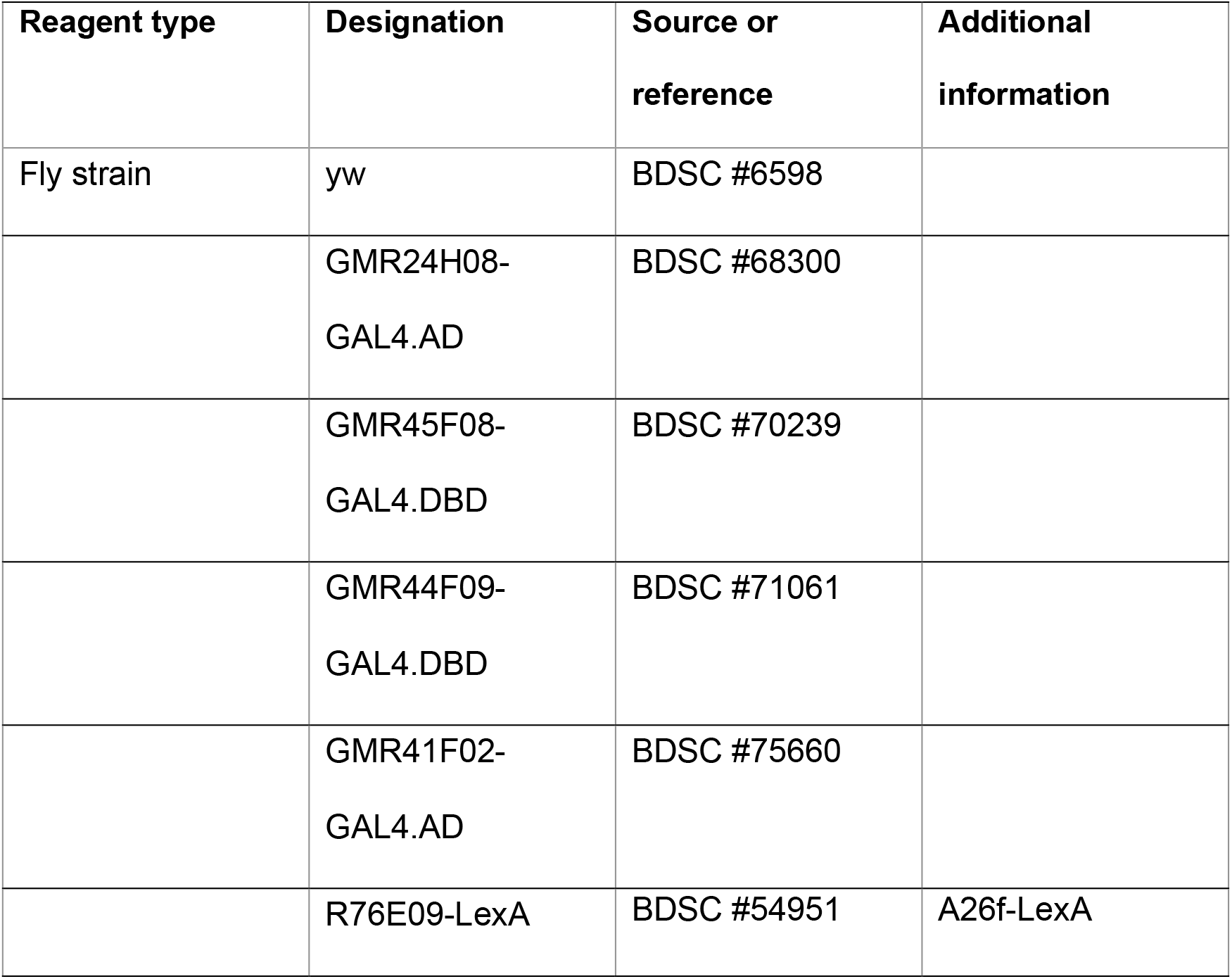
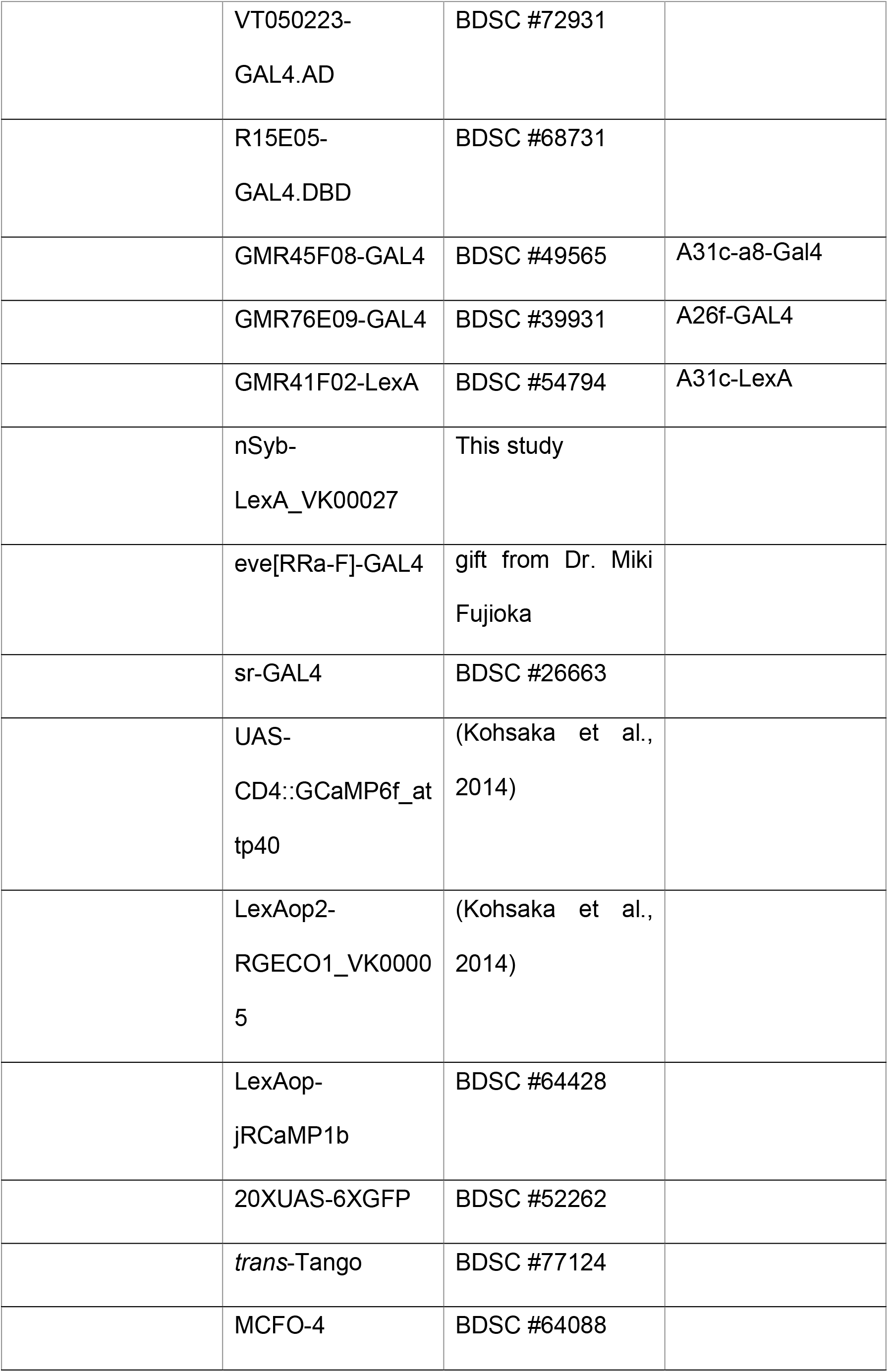
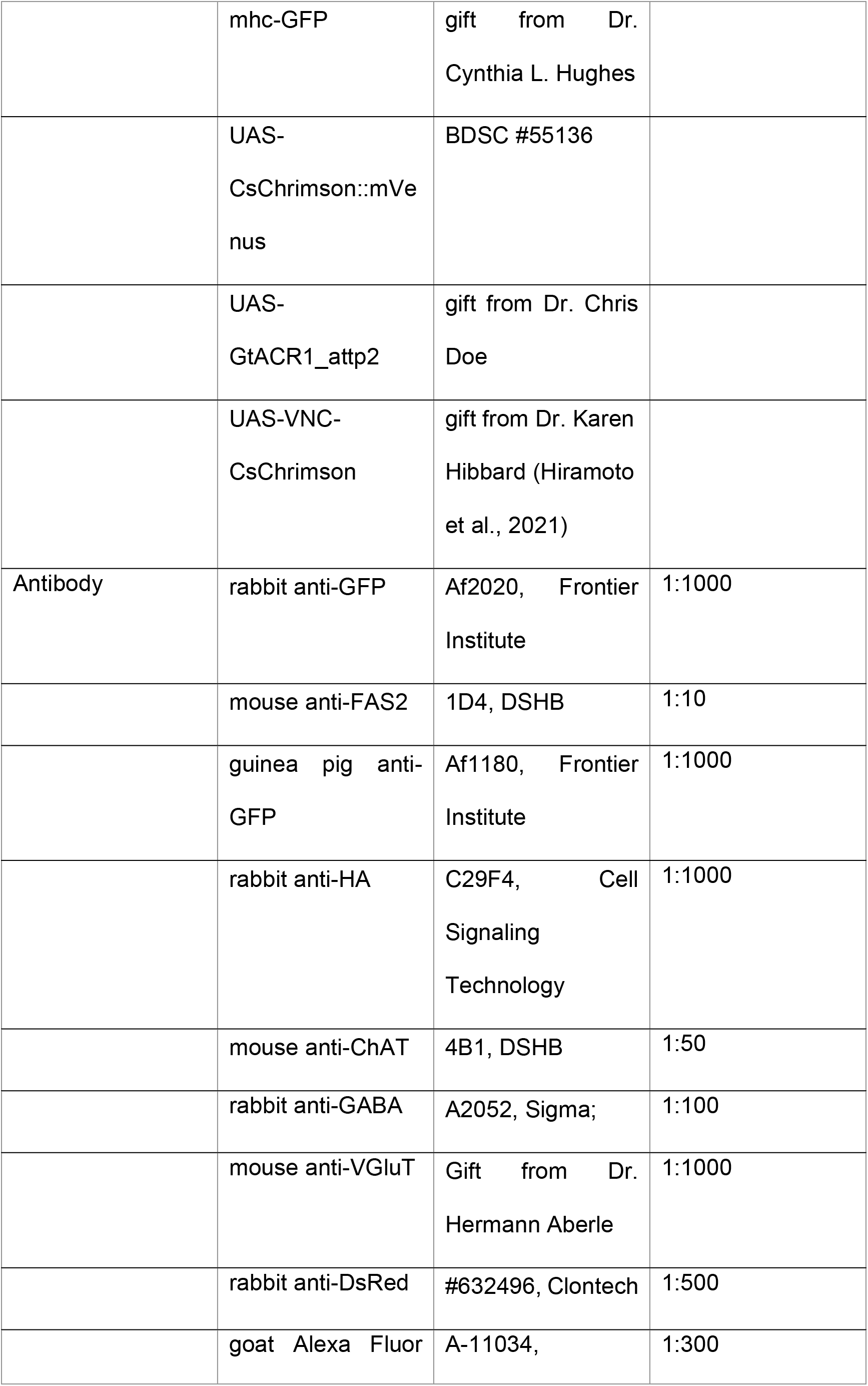
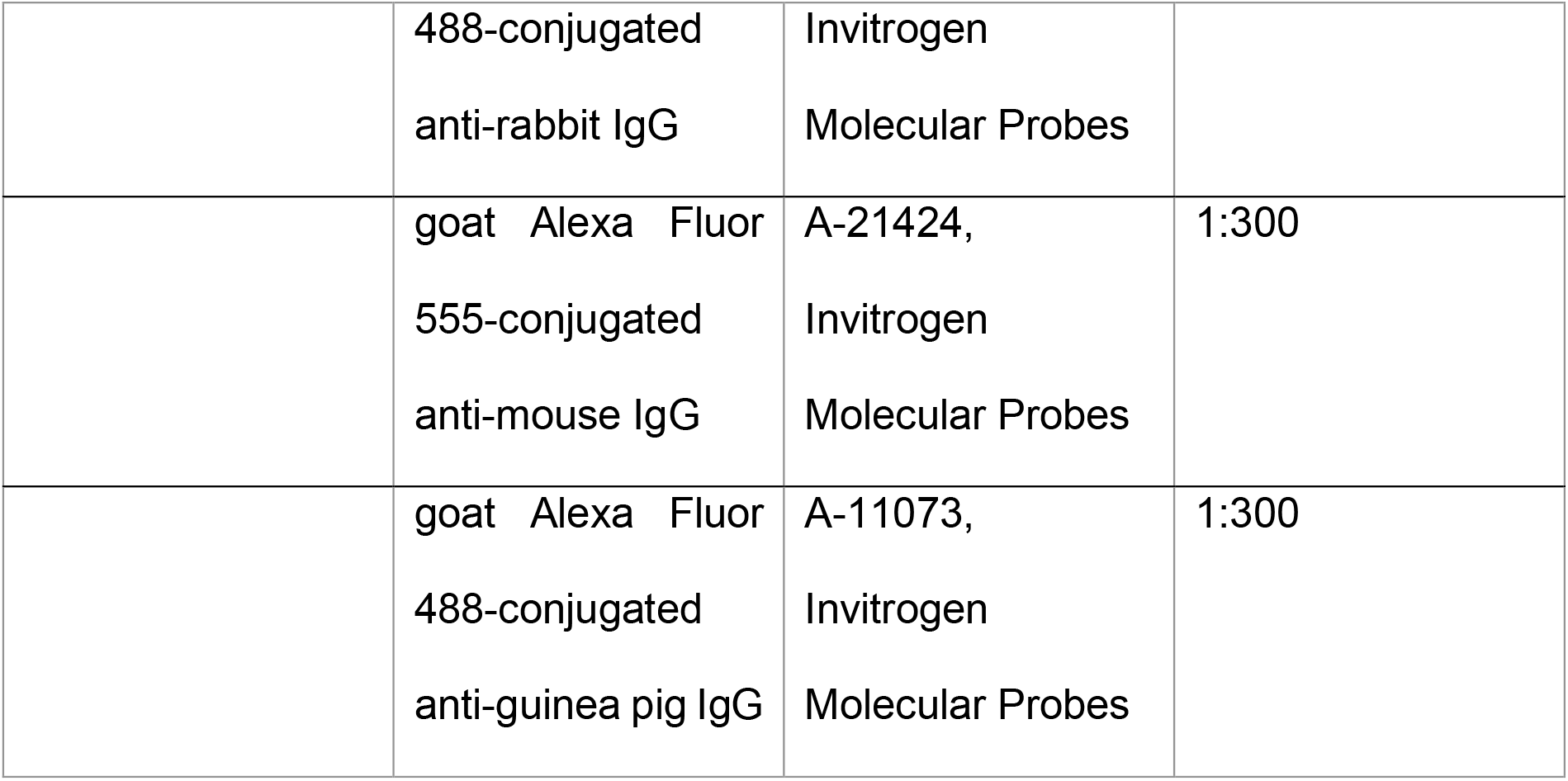
Key resources.

| Reagent type | Designation | Source or reference | Additional information |
| --- | --- | --- | --- |
| Fly strain | yw | BDSC #6598 |  |
|  | GMR24H08-GAL4.AD | BDSC #68300 |  |
|  | GMR45F08-GAL4.DBD | BDSC #70239 |  |
|  | GMR44F09-GAL4.DBD | BDSC #71061 |  |
|  | GMR41F02-GAL4.AD | BDSC #75660 |  |
|  | R76E09-LexA | BDSC #54951 | A26f-LexA |
|  | VT050223-GAL4.AD | BDSC #72931 |  |
|  | R15E05-GAL4.DBD | BDSC #68731 |  |
|  | GMR45F08-GAL4 | BDSC #49565 | A31c-a8-Gal4 |
|  | GMR76E09-GAL4 | BDSC #39931 | A26f-GAL4 |
|  | GMR41F02-LexA | BDSC #54794 | A31c-LexA |
|  | nSyb-LexA_VK00027 | This study |  |
|  | eve[RRa-F]-GAL4 | gift from Dr. Miki Fujioka |  |
|  | sr-GAL4 | BDSC #26663 |  |
|  | UAS-CD4::GCaMP6f_at tp40 | (Kohsaka et al., 2014) |  |
|  | LexAop2-RGECO1_VK00005 | (Kohsaka et al., 2014) |  |
|  | LexAop-jRCaMP1b | BDSC #64428 |  |
|  | 20XUAS-6XGFP | BDSC #52262 |  |
|  | *trans*-Tango | BDSC #77124 |  |
|  | MCFO-4 | BDSC #64088 |  |
|  | mhc-GFP | gift from Dr.<br>Cynthia L. Hughes |  |
|  | UAS-CsChrimson::mVenus | BDSC #55136 |  |
|  | UAS-GtACR1_attp2 | gift from Dr. Chris<br>Doe |  |
|  | UAS-VNC-CsChrimson | gift from Dr. Karen<br>Hibbard (Hiramoto<br>et al., 2021) |  |
| Antibody | rabbit anti-GFP | Af2020, Frontier<br>Institute | 1:1000 |
|  | mouse anti-FAS2 | 1D4, DSHB | 1:10 |
|  | guinea pig anti-GFP | Af1180, Frontier<br>Institute | 1:1000 |
|  | rabbit anti-HA | C29F4, Cell<br>Signaling<br>Technology | 1:1000 |
|  | mouse anti-ChAT | 4B1, DSHB | 1:50 |
|  | rabbit anti-GABA | A2052, Sigma; | 1:100 |
|  | mouse anti-VGluT | Gift from Dr.<br>Hermann Aberle | 1:1000 |
|  | rabbit anti-DsRed | #632496, Clontech | 1:500 |
|  | goat Alexa Fluor | A-11034, | 1:300 |
|  | 488-conjugated<br>anti-rabbit IgG | Invitrogen<br>Molecular Probes |  |
|  | goat Alexa Fluor<br>555-conjugated<br>anti-mouse IgG | A-21424,<br>Invitrogen<br>Molecular Probes | 1:300 |
|  | goat Alexa Fluor<br>488-conjugated<br>anti-guinea pig IgG | A-11073,<br>Invitrogen<br>Molecular Probes | 1:300 |

### Immunostaining and calcium imaging

We used a standard immunostaining procedure (Kohsaka et al., 2014). First, the larvae were dissected in the fillet preparation, fixed in 4% formaldehyde for 30 min at room temperature, washed twice with 0.2% Triton X-100 in PBS (PBT) for 15 min at room temperature, blocked with 5% normal goat serum in PBT for 30 min at room temperature, and stained with the first antibody at 4 °C for 24 to 48 hours. After that, the preparations were washed twice with PBT for 15 min and stained with the second antibody at 4 °C for 24 to 48 hours. Sources and concentrations of antibodies are listed in Table 1.

In the calcium imaging of the isolated CNS, the CNS of third instar larvae was dissected out (Kohsaka et al., 2014), transferred to a drop of TES buffer (TES 5 mM, NaCl 135 mM, KCl 5 mM, MgCl2 4 mM, CaCl2 2 mM, sucrose 36 mM; pH = 7.15), and attached dorsal-up on MAS-coated slide glass for imaging (Matsunami Glass, Japan). GCaMP6f fluorescence was detected by a spinning-disk confocal unit (CSU21, Yokogawa, Japan) and an EMCCD camera (iXon, Andor Technology, Germany) on an upright microscope, Axioskop2 FS (Zeiss, Germany). We used a dual-view system (CSU-DV, Solution Systems, Japan) to perform dual-color calcium imaging for GCaMP and R-GECO1.

### Top-view crawling assay and analysis

Third instar wandering larvae of *sr-GAL4>GFP* (about 0-4 hours after the start of wandering) were used. We transferred a larva onto an agarose plate of a standard concentration (1.5%), waited for about 1 minute, and took a video for 5 minutes. An Olympus stereomicroscope (SZX16, Olympus, Japan) and a 0.7x lens were used for magnification. A CMOS camera (C11440-22CU, Hamamatsu Photonics, Japan) was used for video recording. A square of 1.6 × 1.6 cm of 1024 × 1024 pixels was recorded. The frame rate was set at 30 Hz. A mercury lamp (U-HGLGPS, Olympus, Japan) and an excitation filter (460-495 nm) were used to deliver ~ 5 μW/mm^2^ of blue light for illumination.

We reviewed all videos to extract episodes of straight runs of more than three strides. We then randomly selected three episodes for each larva and analyzed the stride parameters. An ImageJ script was used to manually annotate the video to obtain kinematic parameters (version 1.53, Abràmoff et al., 2004). The stride length was obtained from the distance between the landing positions of the prominent ventral denticle at A8 on one lateral side. The stride duration was obtained from the duration between the unhooking moments. The time of wave initiation was annotated when the A8 prominent denticle moved half a segmental length.

To model the relationship between the stride duration and the duration of the two constituent phases, we tested the polynomial models and the piecewise linear model with two pieces. We then compared the Bayesian information criterion (BIC) between these models (Burnham and Anderson, 2004). The BIC is defined as

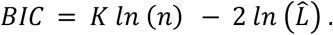

*K* is the number of estimated parameters in the model. *n* is the amount of data. 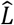 is the maximum value of the likelihood function for the model. In the case of least squares estimation with normally distributed errors, BIC can be expressed as

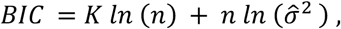

where 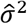 is the average of the squares of residuals. We calculated the BIC for the linear piecewise model of two pieces and the polynomial models of degrees from 2 to 10. The BIC has a minimum value with the cubic polynomial model.

### Side-view imaging of the muscular ends and analysis

Third instar wandering larvae (about 0 to 12 hours after starting wandering) were used. An agarose plate of a standard concentration (1.5%) with black ink (0.2%) was used as the substrate. We oriented a CMOS camera (C11440-22CU, Hamamatsu, Japan) and its zoom lens (MLM3X-MP, Computar, Japan) with a 2x extender (FP-EX2, RICOH, Japan) horizontally for recording. Each time one larva was transferred to the agarose plate for recording. We manually moved the plate to let the camera focus on the larval body wall. The top-view imaging was simultaneously recorded with the same instrument described in the previous method section. A mercury lamp (U-HGLGPS, Olympus, Japan) and an excitation filter (460-495 nm) were used to deliver 5 μW/mm^2^ of blue light for the illumination of the GFP-tagged tendon cells. We recorded at 30 Hz for about 3 minutes and typically collected 3-5 episodes in focus. Each episode includes 2-5 straight crawls.

To analyze the kinematics of the muscular movement, we used DeepLabCut (Mathis et al., 2018) to track the muscular ends. We labeled the muscular ends for 40-50 frames in each video and trained the resnet50 network with the labeled frames for 1,000,000 iterations. To understand the relationship between the contraction of LT muscles and the head and tail movement, an ImageJ script was used to obtain the minimum/maximum length of the LT2 muscle, the maximum thoracic length, the tail traveling distance, and the interwave duration (version 1.53, Abràmoff et al., 2004). To obtain the minimum/maximum length of the LT2 muscle, we annotated the position of the muscular ends of the LT2 muscle in segments A2-A7 when they were mostly contracted and extended and calculated the distance of the pairs of muscular ends. To obtain the maximum thoracic length, we annotated the anterior end of the head and the central point of the T3/A1 segmental boundary at the dorsal side and calculated the distance between them. To obtain the tail traveling distance, we annotated the landing positions of the tail and calculated the distance. The interwave duration was obtained as described in the previous section.

### Trans-synaptic tracing by *trans-*Tango

As *trans*-Tango expression is leaky in larval ventral nerve cord (VNC) neurons when using the recommended rearing temperature 18 °C (Talay et al., 2017), *trans*-Tango larvae were incubated at 30 °C for one day before the experiment. *trans*-Tango expression was thereby restricted to a small number of neurons in combination with the split GAL4 driver *A31c-a8*. We then identified each single neuron by comparing its morphology to the EM database (Ohyama et al., 2015).

### EM reconstruction

Serial sectioning transmission electron microscopy (ssTEM) data were analyzed as described in Ohyama et al., 2015. Briefly, reconstructions were made in a modified version of CATMAID (Saalfeld et al., 2009; http://www.catmaid.org). LT motoneurons and their presynaptic partners had been identified and reconstructed previously within the ssTEM volume (Zwart et al., 2016). These reconstructions were used to identify and reconstruct all presynaptic partners.

### Measurement and quantification of calcium activity

To analyze calcium imaging data, we manually circled regions of interest (ROIs) using ImageJ (version 1.53, Abràmoff et al., 2004). ROIs were chosen at the medial dendritic sites for the A26f neurons, at the axons for the A31c neurons, and the neuropil for the pan-neuronal line in each neuromere. To compare the calcium imaging of different forward cycles, we normalized the time in Figure 2D and Figure 4D relative to the peak ΔF/F of nSyb in segments A4 and A1. We normalized the time in Figure 4F’ relative to the peak ΔF/F of A31c in A4 preceding the FW and the peak ΔF/F of A31c in A4 during the FW. To obtain the time-lagged cross correlation, we slide a trace of calcium activity as in Figure 2D or Figure 4D, calculated the Pearson correlation coefficients with traces of calcium activity in other segments, and calculated the mean value of correlation coefficients by using Fisher-z correction.

### Optogenetic assay of free crawling

We assayed the response of larvae to optogenetic stimulation by using the same imaging system as the top-view imaging assay. The background illumination and the light for the optogenetic stimulation were set as the following. In the GtACR1 groups, we used a 590 nm LED of ~ 150 μW/mm2 to provide the optogenetic stimulation, while a 660 nm LED (M660L3, Thorlabs, USA) or an infrared light (LDQ-150IR2-850, CCS, Japan) provided the background illumination. In the CsChrimson groups, we used an 850 nm infrared light (LDQ-150IR2-850, CCS, Japan) of ~ 40 μW/mm2 to provide the background light and used the 660 nm LED to apply the optogenetic stimulation of ~ 60 μW/mm2. We used an ImageJ script to manually annotate videos to obtain the kinematic parameters (version 1.53, Abràmoff et al., 2004). In the experiments using GtACR1, the larva can show transient turning or stopping responses to 590 nm light. In these groups, we analyzed strides if forward cycles were not halted or after forward cycles were reinitiated. In the experiment using *A26f-sp* drivers, we only analyzed the data when the *GtACR1/CsChrimson* was expressed in more than four A26f neurons, which was determined by post-hoc staining.

### Assay of muscular response to optogenetic stimulation in the fillet and sideways preparation

We used a semi-intact fillet preparation to assay muscular responses to optogenetic activation (Kohsaka et al., 2014). After the preparation, we waited for about 10 minutes, until the larva stopped its frequent spontaneous axial waves.

To constrain the movement of the larva without impairing peristaltic behavior and visualize the lateral side of the larva, we devised a new preparation named sideways preparation. In this preparation, the larva is fixed by two pins on a vertical side of a Polydimethylsiloxane (PDMS; Silpot 184, Toray, Japan) plate and oriented lateral side up to visualize the LT muscles. The larva can show spontaneous forward peristalsis-like behavior in this preparation. In the preparation, we prepared a PDMS plate with a standing PDMS island filled with 4°C TES buffer, transferred a larva to the PDMS plate, and used two pins to fix the head and tail of the third instar larvae (Figure 5A and 5B). The tail was pinned to the bottom PDMS substrate to make the pin perpendicular to the larval sagittal plane with two pricking points close to the two prominent lateral denticles in the A8 segment. The head was pinned to the PDMS island to make the pin perpendicular to the larval frontal plane. After pinning, the PDMS island was attached to the tail pin and supported the ventral larval body. 4°C TES buffer was used to reduce the larval motion during the preparation. We changed the buffer to 25 °C before imaging.

A local stimulation microscope was used for muscular imaging and optogenetic stimulation (Matsunaga et al., 2013; Takagi et al., 2017). The microscope (FV1000, Olympus, Japan) has two separate optical paths for muscular imaging and optical stimulation, respectively: blue light from a Xeon lamp (X-Cite exacte, Excelitas Technologies, US) and a GFP dichroic mirror (U-MGFP/XL, Olympus, Japan), which were used to image the muscles in the abdominal segments A3/A4 to A7/A8, and a scanning laser of blue (488 nm) or green (559 nm) light, which was used to stimulate the CNS optogenetically. A dichroic mirror separates the two optical paths. To fit the larva into the field of view, we used a 4x Olympus objective and a 1 x or a 0.63 x adapter. Muscular contractions were recorded by an EMCCD camera (iXon, Andor Technology, Germany). We used different combinations of optogenetic stimulation and muscular illumination. In the sideways preparation, a rectangular scanning of about 0.85 mm x 0.4 mm by the 559nm laser was used for optogenetic stimulation (~ 20 μW/mm2 for the CsChrimson groups and ~ 40 μW/mm2 for the GtACR1 groups), while blue light of ~ 10 μW/mm2 was used for muscular illumination. In the fillet preparation, a spiral scanning of a radius of ~ 0.3mm by a 488 nm laser was used to activate the Chr2 (~ 400 mW/mm2), while a blue light of ~ 50 μW/mm2 was used for muscular illumination. DeepLabCut (Mathis et al., 2018) was used to track the muscular ends. We labeled the muscular ends in 40-50 frames in each video and trained them using the resnet50 network. The neural network was trained 1,000,000 times.

### Statistical tests for the optogenetic experiments

Changes in muscle length were normalized to the minimal muscle length within each animal. Changes in other values (stride duration, stride length, etc.) were directly used for statistical tests. We tested the significance of the changes before and after the optogenetic manipulation and compared the changes between the experimental and the control. As each animal was treated with optogenetic stimulation multiple times, to increase the statistical power and to avoid Type-I error (false positive), we used hierarchical bootstrapping methods for the comparison before and after the optogenetic stimulation and the comparison between the experimental and control group (Saravanan et al., 2020). To generate the bootstrapped dataset, we resampled data from the experimental dataset 10,000 times. Each time we 1) resample *n* animals with replacement (*n* is the animal number in the experiment), and 2) resample *m*_1_,.., *m*_*n*_ trials within animals with replacement (*m*_*i*_ is the number of trials of the resampled animal *i* in the experiment) (Saravanan et al., 2020). For the comparison before and after the optogenetic stimulation, we used the empirical method by 1) computing the animal-wide mean of the bootstrapped sample *μ*^*^, 2) computing the difference between *μ*^*^ and the animal-wide mean of the experiment *μ*, and 3) computing the p-value as the quantile of *μ* in *μ*^*^ – *μ* (Efron and Tibshirani, 1994). For the comparison between the experimental and control group, we 1) computed the animal-wide mean of the two bootstrapped samples (*μ*^*a*,*^ and *μ*^*b*,*^), 2) computed a joint probability distribution of *μ*^*a*,*^ and *μ*^*b*,*^, and 3) computed the p-value as the density of the joint probability (Saravanan et al., 2020). All analysis was done with Python (version 3.9.12) scripts using the libraries NumPy (version 1.21.5) and SciPy (version 1.7.3). Asterisks represent the range of p values (*p<0.05; **p<0.005; ***p<0.0005).

## Acknowledgment

We thank Bloomington *Drosophila* Stock Center, KYOTO *Drosophila* Stock Center, and Drs. Chris Doe, Miki Fujioka, Karen Hibbard, and Cynthia L. Hughes for the fly lines (Table 1). We thank Developmental Studies Hybridoma Bank and Dr. Hermann Aberle for the antibodies. We thank Drs. Stefan Pulver and Wenchang Li for critical comments on the paper. This work was supported by MEXT/JSPS KAKENHI grants (17K19439, 19H04742, 20H05048 to A.N. and 17K07042, 20K06908 to H.K.) and the Royal Society of Edinburgh (grant 64553 to M.F.Z.).

## Availability of data and materials

The data generated in this study will be available from the Zenodo repository.

---

**Figure 1 – supplement.**
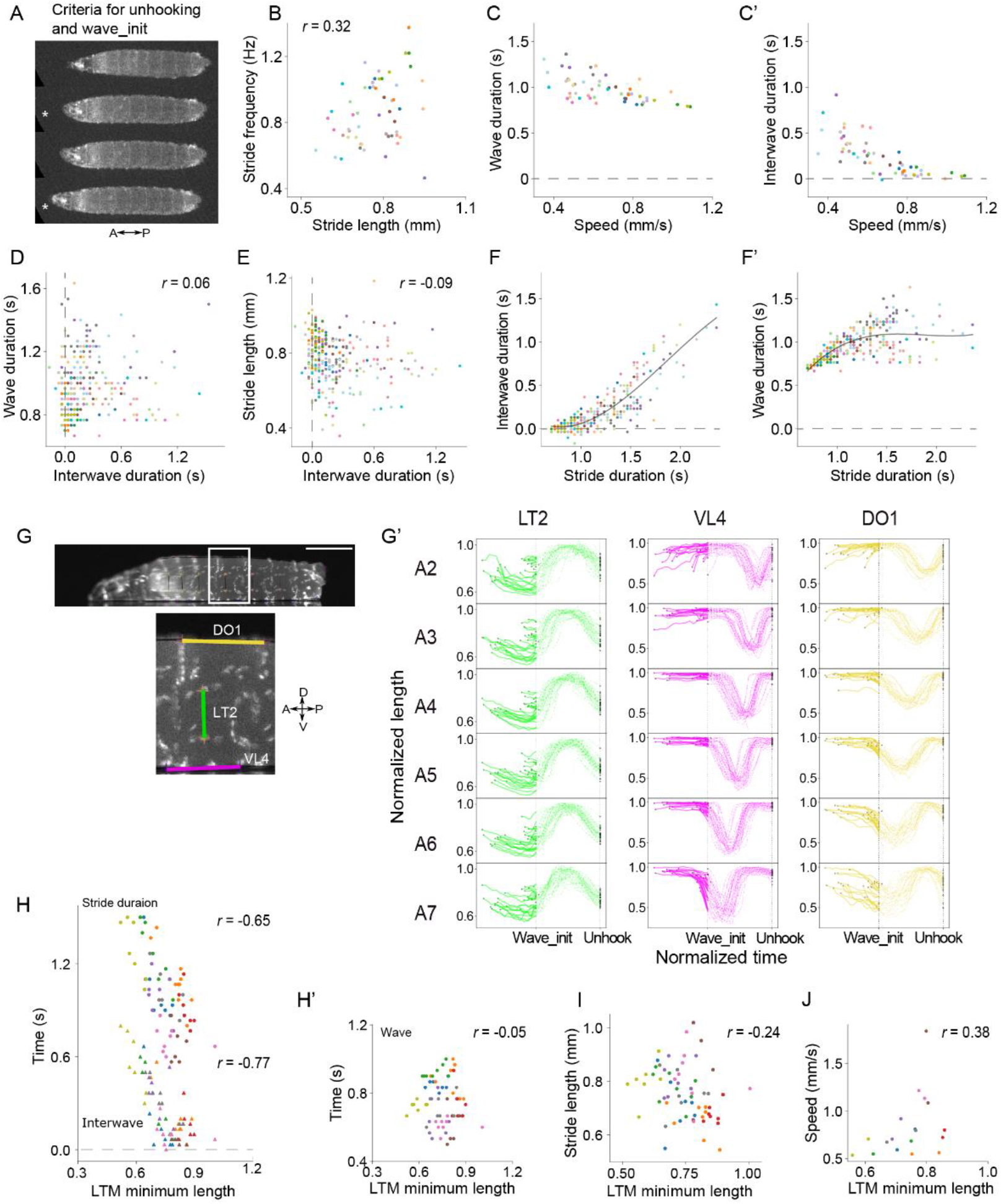
**(A-F)** Supplement to top-view crawling assay. **(A)** Criteria to determine the unhooking moment and the wave initiation moment (See Materials and methods for details.) Asterisks indicate the key frames (upper asterisk: unhooking moment; lower asterisk: wave initiation moment). **(B)** Relationship between stride frequency and stride length. **(C-C’)** Relationship between speed and the two component phases (Pearson correlation coefficient: −0.74 for interwave phase and −0.62 for the wave duration; p-value = 0.127). **(D)** Relationship between the duration of wave phase and interwave phase. **(E)** Relationship between stride length and the duration of interwave phase. **(F)** Relationship between stride duration and the duration of component phases. Lines: cubic regression (regression coefficients: interwave, [0.78, −2.1, 1.7, 0.3]; wave, [0.78, 3.1, −1.7, −0.3]; See Materials and methods for details.) **(G-J)** Supplement to side-view crawling assay. **(G)** An example frame of side-view recording. Scale bar: 1 mm. Lower panel is an enlarged image of the region indicated by the white rectangular in the top panel. The range of muscles (DO1, LT2, and VL4) is labeled. **(G’)** Dynamics of muscle length during forward crawling measured from the side-view assay. Muscle length is normalized to 0-1 by 0 and the maximum length of the muscle. Time is aligned by the wave_init moment and the unhooking moment (See Materials and methods for details). Each trace initiates and ends at the beginning and end of a locomotor cycle.

**Figure 2-supplement.**
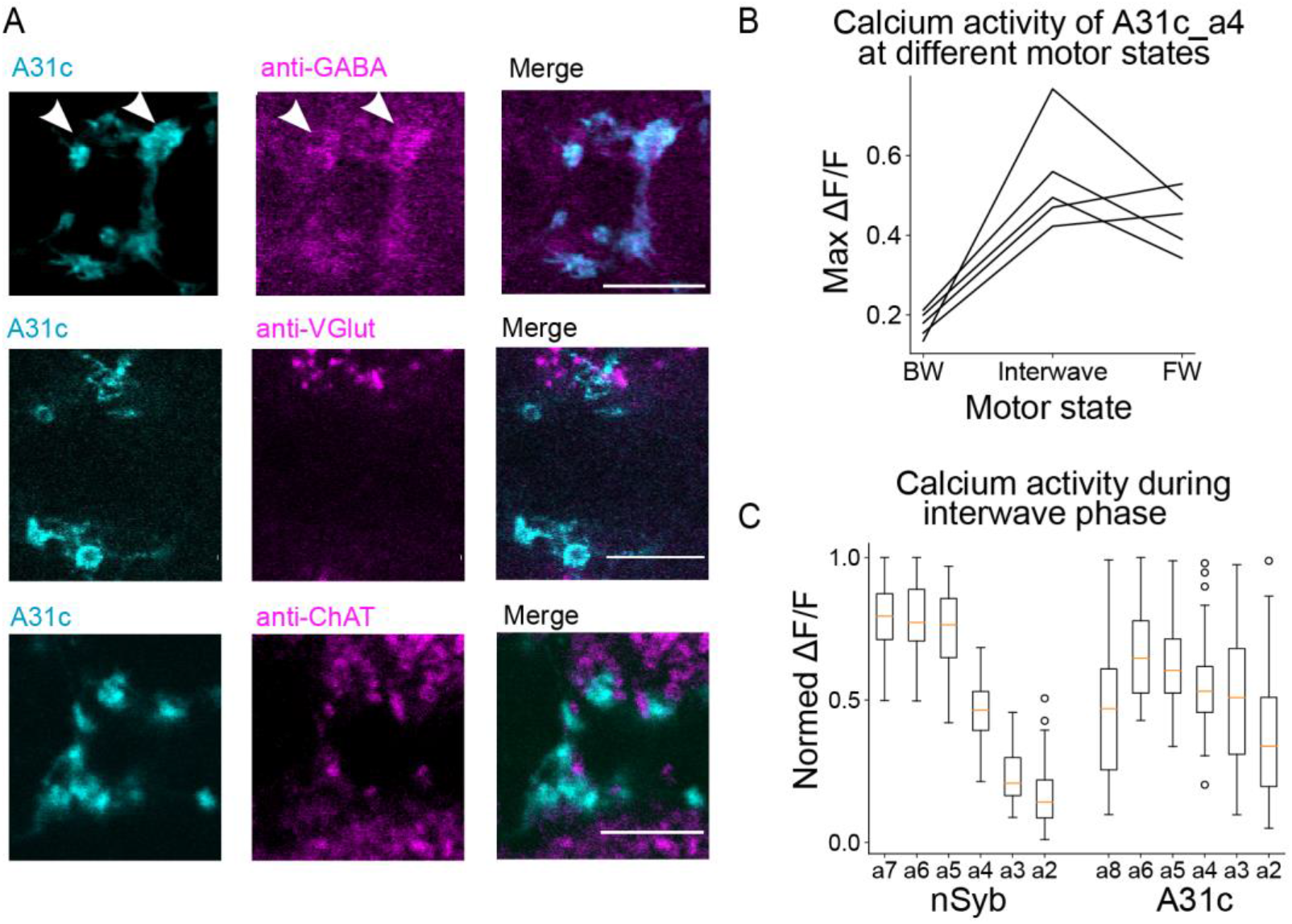
**(A)** Immunostaining reveals that A31c neuron is GABAergic. Scale bars: 10 μm. **(B)** Comparison of the amplitude of the peak calcium signals of the A31c_a4 neuron in different motor states (BW: backward wave; PSync: posterior synchronous burst; FW: forward wave). **(C)** Comparison of the amplitude of the peak calcium signals of nSyb neurons and A31c neurons when nSyb-a7 reaches the peak (See magenta arrowheads in Figure 2E.)

**Figure 3-supplement.**
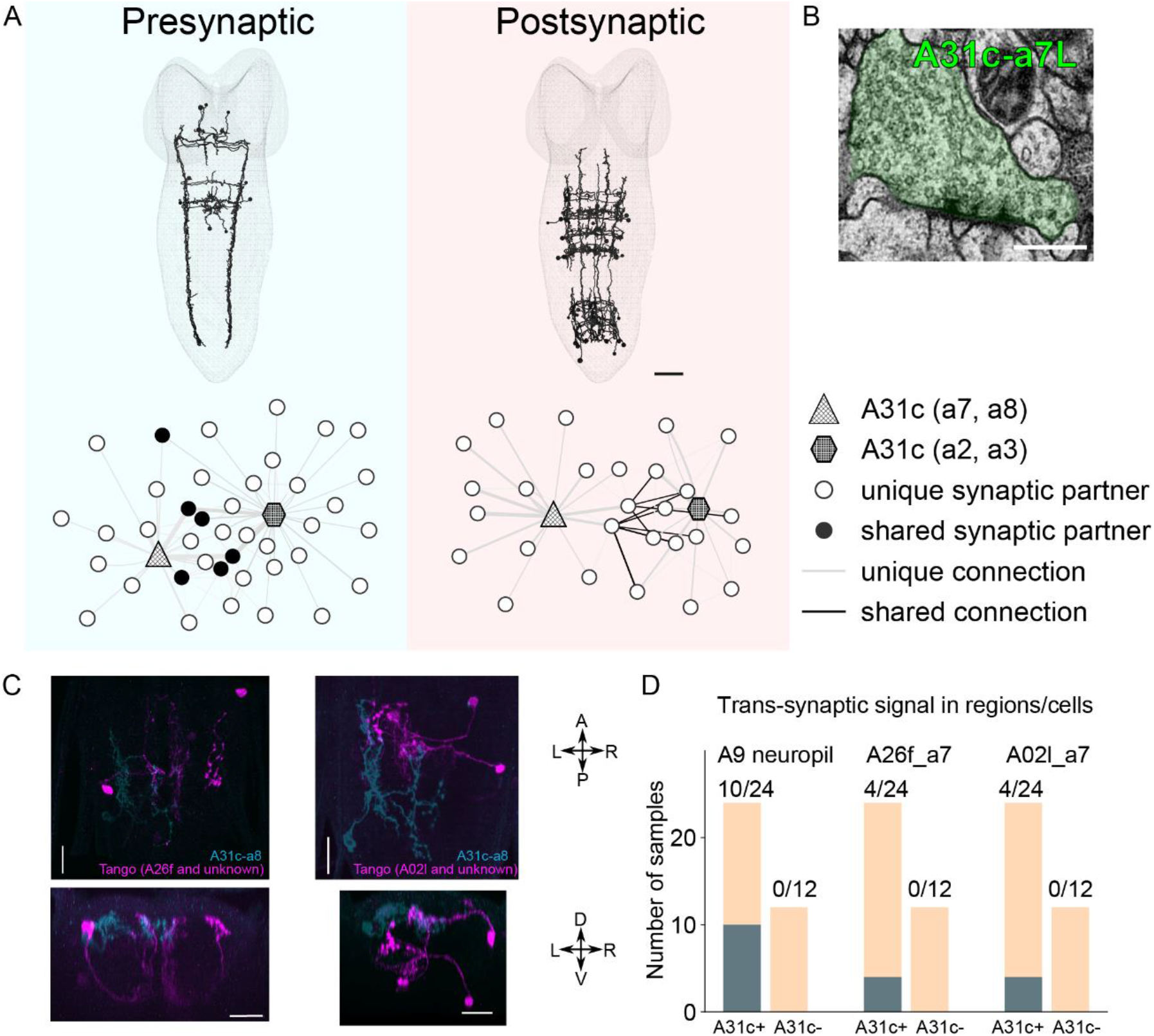
**(A)** Reconstructed main synaptic partners of A31c neurons. **(B)** A sample EM image shows the A31c presynaptic site. **(C)** *trans-*Tango reveals the postsynaptic partners (magenta) of A31c neurons (cyan) (*A31c-a8-sp>trans-Tango*). **(D)** Quantification of the number of samples that shows expression in regions or neurons repeatedly targeted by *trans*-Tango. Scale bars: 20 μm.

**Figure 4-supplement.**
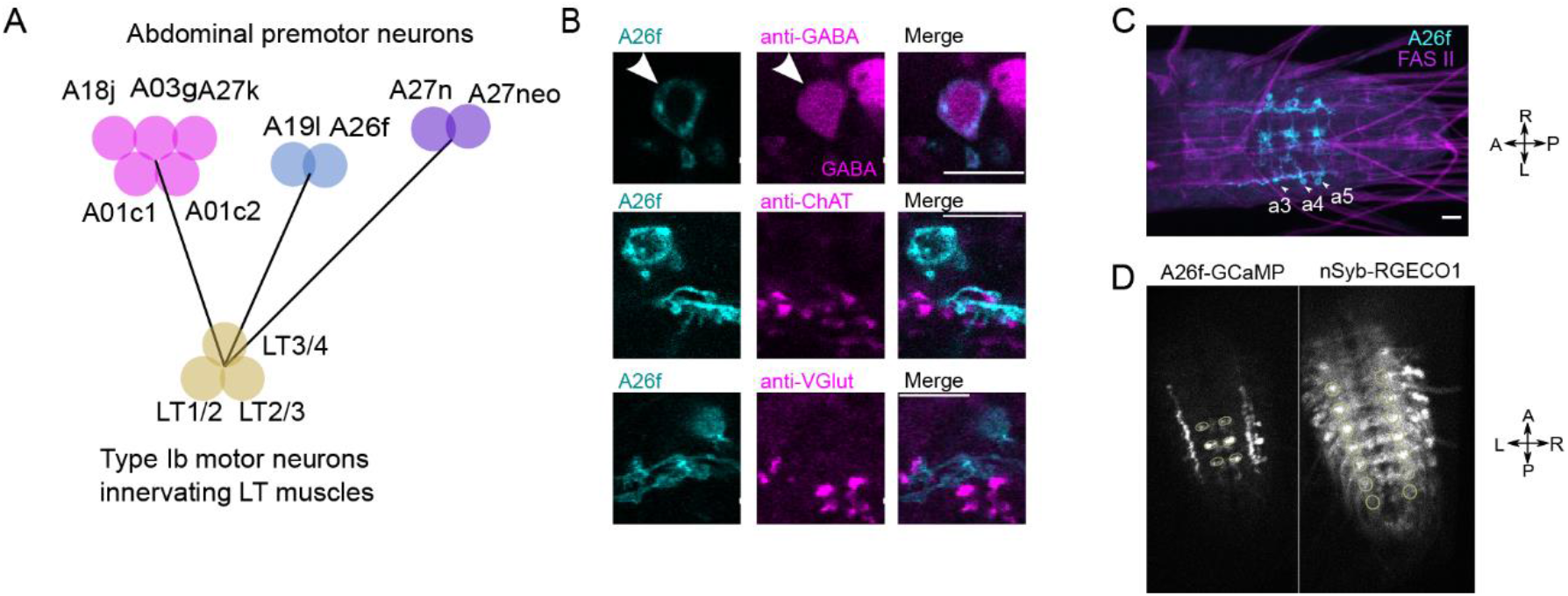
**(A)** Premotor interneurons that target all the motor neurons innervating LT muscles. **(B)** Immunostaining reveals that A26f neuron is GABAergic. Scale bars: 10 μm. **(C)** *A26f-sp* labels A26f neurons in A3 to A5 neuromeres. **(D)** Calcium signals of A26f neurons and nSyb neurons in fictive locomotor cycles.

**Figure 5-supplement.**
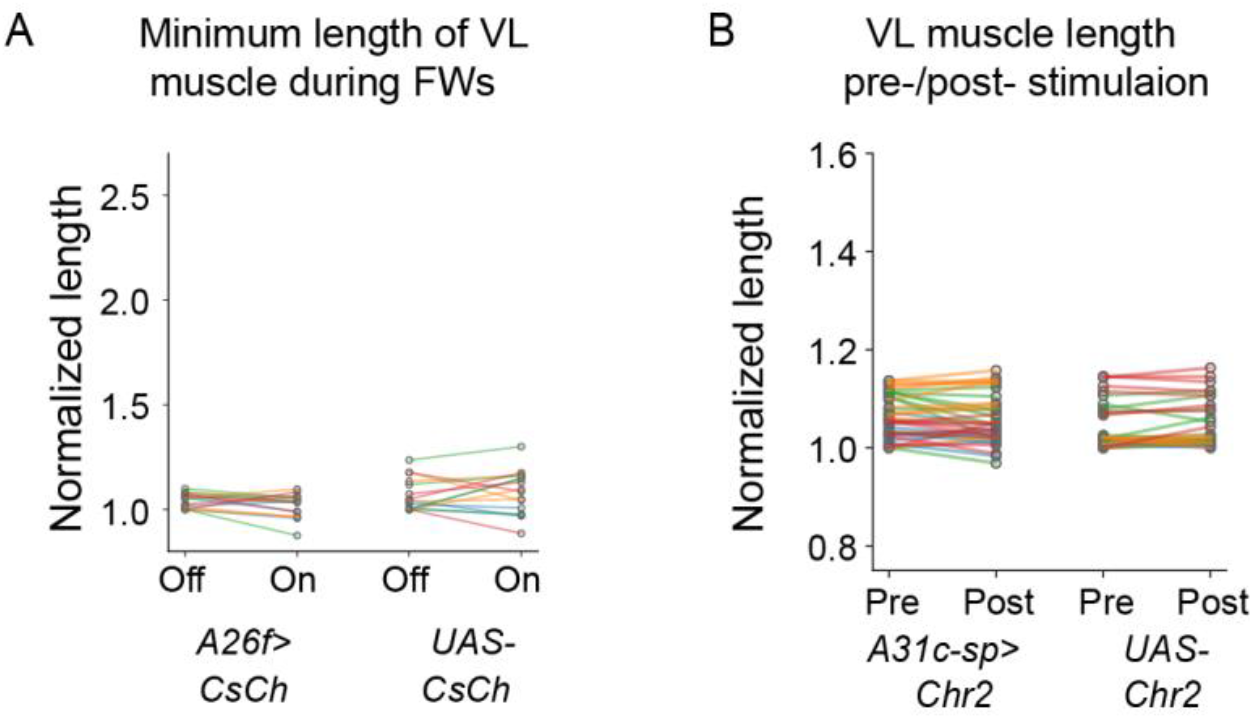
**(A)** The minimum length of longitudinal VL muscles was not affected by optogenetic activation of A26f neurons. **(B)** The length of VL muscles in the resting state was not affected by optogenetic activation of A31c neurons.

**Figure 7-supplement.**
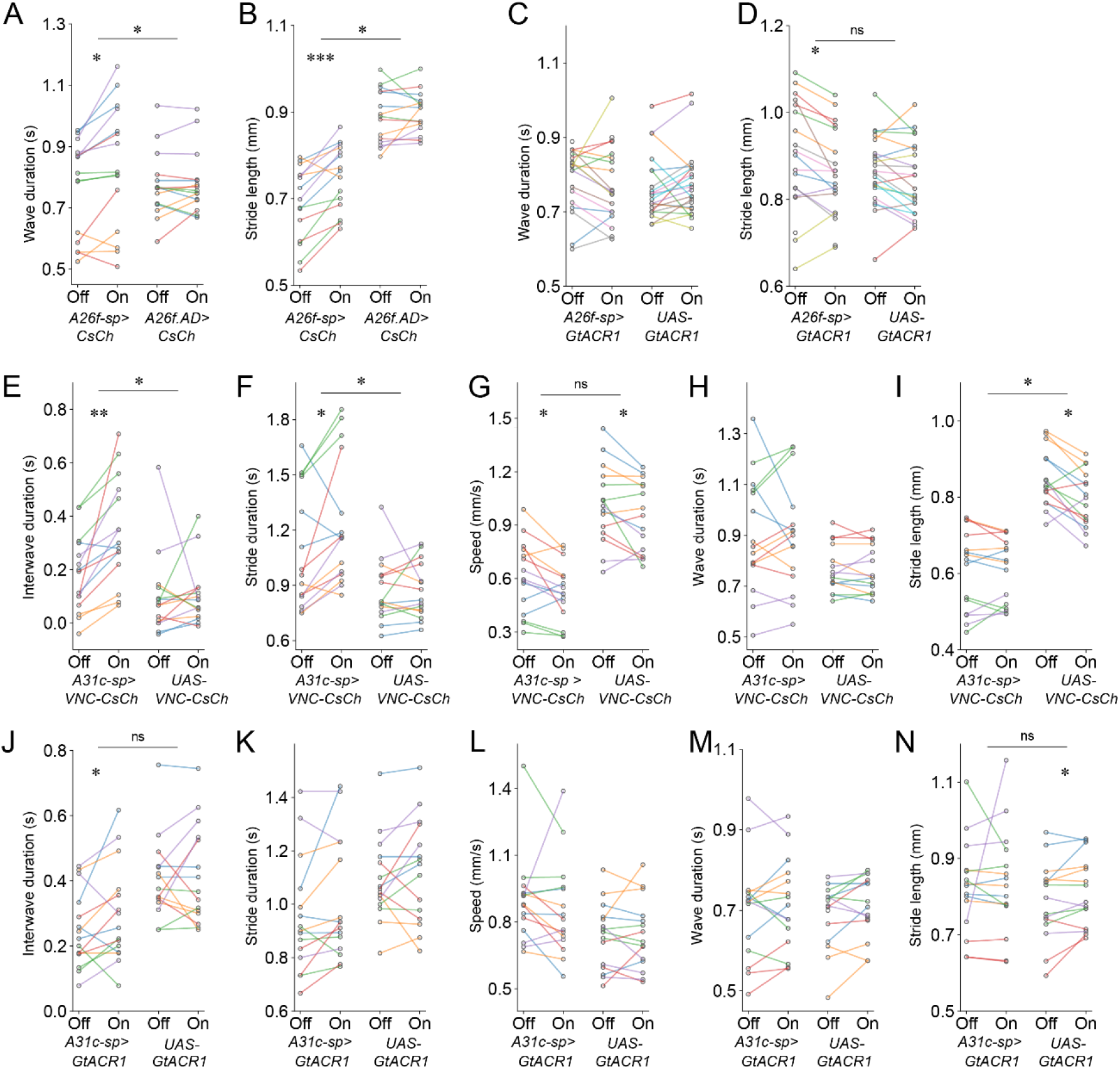
Analysis of crawling kinematics of larvae with optogenetic perturbation. **(A and B)** Activation of A26f neurons. **(C and D)** Inhibition of A26f neurons. **(E-I)** Activation of A31c neurons. **(J-N)** Inhibition of A31c neurons. Statistical results are obtained by the hierarchical bootstrap test (See Materials and methods for details.)

